# Random mtDNA mutagenesis maps genotype-phenotype associations across metabolic contexts and lineages

**DOI:** 10.64898/2026.10.08.757705

**Authors:** Amir Giladi, Ron Baik, Chih-Yao Chung, Minsoo Kim, Josephine Adams, Rahul Saha, Max Land, Ruobing Cui, Emily DeBitetto, Megan E. Kelley, Craig B. Thompson, Ed Reznik, Roshan Sharma, Ronan Challinge, R. Stefan Isaac, Josi Peterson, Alexander van Oudenaarden, Agnel Sfeir

## Abstract

Mitochondrial DNA (mtDNA) mutations cause inherited disease, accumulate with age, and recur in tumors, yet their effects remain unresolved because existing models interrogate one engineered allele at a time. We developed mtDIVE, a random C→T mutagenesis platform for pooled mitochondrial forward genetics in human epithelial cells and zebrafish embryos. mtDIVE generates genotypes ranging from single substitutions to >100 mutations per molecule, impairs respiration, and causes lineage-specific developmental delay. Coupling single-cell transcriptomics with mtDNA genotyping revealed that selection on mutant genomes and their transcriptional consequences depend on metabolic state and developmental lineage. Obligate OXPHOS eliminates highly mutated cells by apoptosis, whereas glycolytic conditions promote their competitive displacement. Individual loci exert modest but non-redundant effects that predict cell state beyond aggregate mutation burden, and cancer-recurrent variants are enriched among mutations linked to reduced mitochondrial transcription. Thus, mtDIVE links mtDNA genotypes to cellular phenotypes, separating the consequences of mutation identity from mutational load.

## INTRODUCTION

Mitochondria are essential multifunctional organelles responsible for cellular energy production and numerous biosynthetic and signaling processes. Human mitochondrial function relies on a multi-copy, 16.6 kb circular mitochondrial DNA (mtDNA) that encodes 13 subunits of the oxidative phosphorylation (OXPHOS) system together with the rRNAs and tRNAs required for mitochondrial translation. Proper mitochondrial function depends on tight coordination between the small mtDNA genome and more than a thousand nuclear-encoded proteins, placing mtDNA integrity at the center of cellular homeostasis^1^.

Since the discovery of mtDNA in the 1960s, more than 250 pathogenic mtDNA mutations have been identified in humans, collectively affecting approximately 1 in 5,000 individuals^2–5^. These disorders exhibit striking variability in clinical presentation and disease progression, even among patients carrying the same mutation^6^. This heterogeneity arises in part from heteroplasmy, the coexistence of wild-type and mutant mtDNA alleles within a cell or a tissue, and in part due to the interplay among variant identity, mutational load, and cellular and tissue contexts^2,7^. In addition to germline variants, low-abundance mtDNA mutations arise somatically and accumulate with age, generating mosaic patterns of mitochondrial genetic variation across cells and tissues that have been linked to aging and age-related disease, including metabolic disorders and neurodegeneration, and more recently associated with cancer^2,3,8–18^. Whether such low heteroplasmy mutations causally influence cellular physiology, or merely mark cells that are already dysfunctional has remained difficult to resolve.

Several systems have been developed to model mtDNA mutations, including transmitochondrial cytoplasmic hybrid cells (cybrids)^19^, proofreading-deficient POLG mutator mice^20,21^, and targeted mtDNA editing with TALE- or zinc-finger-based base editors fused to the bacterial split DddA cytidine deaminase (split-DdCBE) or the TadA-derived adenine deaminases^22–27^. Although valuable, these approaches have significant limitations in mapping the cellular effects of mtDNA variants. POLG-driven mutagenesis is confounded by indels, large deletions, and mtDNA depletion, so a phenotype cannot be attributed to substitutions alone. Targeted editors, by contrast, require a dedicated protein design per locus, hindering high-throughput interrogation. Single-cell mitochondrial profiling now resolves complex mitochondrial genotypes alongside chromatin and transcriptional state^28–32^, so the measurement has outpaced the perturbation, as no system that introduces random, combinatorial mtDNA mutations at controlled heteroplasmy has been developed thus far. Such a platform, deployable across human cells and developing organisms, would allow the functional landscape of mtDNA variants to be mapped rather than sampled one allele at a time. This is especially true in development, where embryos must meet their energetic and biosynthetic demands before the organs of metabolic homeostasis exist; in teleosts, the extraembryonic yolk syncytial layer supplies glucose by converting yolk-derived glutamate through mitochondrial phosphoenolpyruvate carboxykinase (PEPCK-M, *pck2*), an enzyme utilizing GTP from substrate-level phosphorylation in the Krebs cycle rather than by oxidative phosphorylation^33^. How far early vertebrate development can proceed on such OXPHOS-independent routes to ATP and glucose has not been testable with a mutagenesis system that titrates mtDNA damage across a whole embryo and reads genotype together with cell state.

We developed a random mtDNA mutagenesis platform, **mt**DNA **D**iversification **I**n **V**ivo via **E**diting (mtDIVE), that couples the DddA cytidine deaminase to the mtDNA-packaging factor TFAM and is deployed in two formats: 1) an inducible split-DddA system for time- and dose-controlled mutagenesis in non-transformed human retinal pigment epithelial (RPE) cells and 2) an RNA-encoded single-fusion construct for synchronous mutagenesis in zebrafish embryos. Transient mutagenesis impaired respiration and reduced proliferative capacity in both systems, and caused lineage-specific developmental delay in embryos. Single-cell multi-omic profiling showed that mtDIVE activity, the selection acting on mutant genomes, and the transcriptional response to them all varied with human metabolic context and zebrafish cell lineage. Finally, the identity of the mutated loci predicted cellular state beyond mutation burden alone, and this functional axis was occupied preferentially by mtDNA mutations that recur in human tumors. Together, mtDIVE links genotype, mutational load, metabolism, and lineage to cellular fate within a single experimental framework.

## RESULTS

### mtDIVE introduced random C→T substitutions across the mitochondrial genome of human cells and zebrafish embryos

To systematically dissect the functional consequences of mtDNA mutations, we developed a random mutagenesis platform, termed mtDNA Diversification In Vivo via Editing (mtDIVE), designed to introduce base substitutions across the mitochondrial genome at varying heteroplasmy. The core effector fuses a mitochondrial localization sequence (MLS), the mtDNA-binding factor TFAM (HMG-A and HMG-B domains), a DddA cytidine deaminase, and a uracil glycosylase inhibitor (UGI) (Fig. 1A). Unlike TALEN-based editors, TFAM tethers to mtDNA in a non-specific manner, where DddA converts neighboring cytidine to uridine, and UGI blocks uracil excision so that edits are fixed as C→T (G→A) substitutions^23^. We implemented two delivery strategies. For *in vivo* use, we transcribed mRNA encoding the full construct and injected it into one-cell-stage zebrafish embryos (zmtDIVE, Fig. 1B), producing a transient pulse of mtDNA mutagenesis during early development. For *in vitro* use in human cells, we generated a stably integrated, inducible construct in human retinal pigment epithelial (RPE) cells (hmtDIVE, Fig. 1C). Here DddA was split across the chemically inducible dimerization domains FRB and FKBP, which associate only in the presence of rapalog (AP21967)^34,35^. The TFAM-FKBP-DddA half was additionally placed under a doxycycline-inducible promoter. This dual control allows expression (doxycycline) and reconstitution of active deaminase (rapalog) to be gated independently (Fig. 1C). Immunofluorescence (IF), subcellular fractionation, and western blotting confirmed proper mitochondrial localization and inducible expression of both halves (Fig. 1D, S1A-C). We then tested the capacity of the system to generate mtDNA mutations, using bulk sequencing of human samples and of whole zebrafish embryos (Methods). Both zmtDIVE and hmtDIVE induced mutations throughout the mitochondrial genome, generating a spectrum dominated by TC→TT substitutions, consistent with the 5’-TC-3’ dinucleotide substrate preference of DddA (Fig. 1E-G, S1D)^23^.

**Figure 1.**
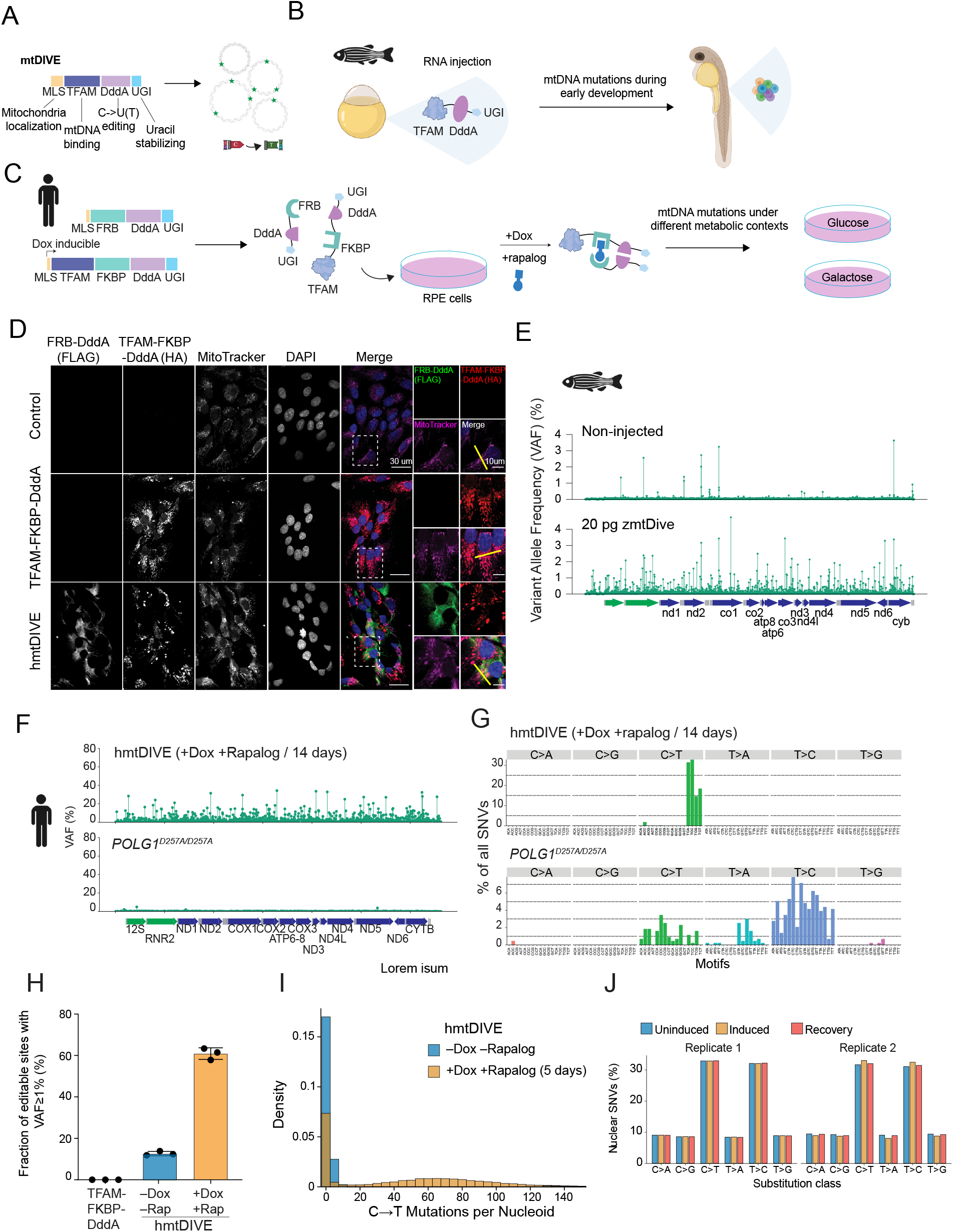
mtDIVE, an inducible and tunable platform for generating mtDNA mutations in cultured human cells and zebrafish embryos. **(A)** Schematic of the core mtDIVE construct. A mitochondrial localization signal (MLS) targets the fusion protein to mitochondria, where TFAM binds mtDNA, positioning the double-stranded DNA-specific cytidine deaminase DddA to catalyze C→U editing; the uracil glycosylase inhibitor (UGI) inhibits uracil excision, allowing mutations to fix as C→T substitutions. Expression of this construct results in cells harboring random de novo mtDNA mutations (green dots). **(B)** Schematic of zmtDIVE-induced mutagenesis in zebrafish embryos. RNA encoding the MLS-TFAM-DddA-UGI fusion was injected into one-cell-stage embryos, inducing a transient pulse of mtDNA mutagenesis during early embryogenesis. **(C)** Schematic of the doxycycline- and rapalog-inducible split hmtDIVE system in human RPE cells. DddA is split into two halves fused to inducible dimerization domains FRB and FKBP; the TFAM-FKBP-DddA fragment is additionally placed under a doxycycline-inducible promoter. Rapalog drives FRB–FKBP heterodimerization and reconstitution of active DddA. Following induction, cells were cultured in glucose- or galactose-containing medium. **(D)** Representative immunofluorescence images of RPE cells after 48 hours of doxycycline and rapalog treatment. WT, TFAM-FKBP-DddA, and hmtDIVE cells were stained for HA-tagged TFAM–FKBP–DddA and FLAG-tagged FRB–DddA. Mitochondria were visualized with MitoTracker, and nuclei were stained with DAPI. Yellow lines indicate the regions used for line-scan analysis in Fig. S1A. Scale bar, 30 μm. Insets display zoom-in regions; scale bar 10 μm. **(E)** Lollipop plots showing bulk mtDNA variant allele frequencies (VAFs) in non-injected (top) and 20 pg zmtDIVE-injected zebrafish embryos at 24 hours post fertilization. Data are from a representative replicate from two profiled embryos (n=2). **(F)** Lollipop plots showing bulk mtDNA VAF in hmtDIVE-expressing RPE cells after 14 days of doxycycline (Dox) and rapalog treatment. Cells expressing proofreading-deficient POLG^D257A/D257A^ were cultured for over 3 months and analyzed for comparison. Data are representative from (n=4) biological replicates. (E-F) Bottom mitochondrial genome maps position detected variants relative to mtDNA-encoded genes. Protein-coding genes are shown in blue, rRNA in green and tRNA genes in gray. **(G)** Mutational signature analysis of mtDNA variants detected in hmtDIVE-expressing cells after 14 days induction and in POLG^D257A/D257A^ cells. Histograms show single-nucleotide variant (SNV) frequencies according to trinucleotide sequence context. Data are representative of n=4 biological replicates. **(H)** Fraction of mtDNA editable TC motifs with VAF >1% in TFAM-FKBP-DddA controls and hmtDIVE-expressing cells with or without induction. Cells were grown for 5 days. *n* = 3 biological replicates per condition; Data are presented as mean ± SD. **(I)** Density plots showing the number of detected mtDNA mismatches per MtFiber-seq full-length mtDNA molecule derived from hmtDIVE-expressing RPE cells with or without Dox and rapalog induction. **(J)** Distribution of nuclear single nucleotide (SNV) variants detected by MtFiber-seq long-read sequencing in hmtDIVE-expressing RPE cells across three conditions: uninduced, after 5 days induction, and following 21 days drug withdrawal (recovery). SNVs are grouped by substitution class. Two biological replicates are shown.

In zebrafish, zmtDIVE mtDNA editing showed reproducible patterns across mtDNA loci that scaled with injection dose (Fig. 1E, S1E), whereas control constructs lacking TFAM or carrying a catalytically inactive DddA domain showed no detectable mutagenic activity (Fig. S1F). In RPE cells, 14 days of hmtDIVE induction produced hundreds of heteroplasmic variants (variant allele frequency [VAF] > 0.1%) distributed across the mitochondrial genome (Fig. 1F). Rapalog titration identified 50 nM as the optimal dose that balances cell viability (Fig. S1G) against editing efficiency (Fig. S1H), and this concentration was used throughout. Active editing of mtDNA by hmtDIVE resulted in a modest reduction in mitochondrial branch length, an effect exacerbated when glucose was withdrawn (Fig. S1I-J). Compared with cells expressing POLG1-D257A^8,20^, which lack proofreading activity, hmtDIVE introduced substantially more variants at higher VAF within the TC sequence context (Fig. 1F-G). After 5 days of induction, ∼60% of all editable 5′-TC-3′ motifs in the genome exhibited VAF ≥ 1% (Fig. 1H). Editing was tightly controlled, as Dox induction drove a marked accumulation of variants, with minimal uninduced leakiness (Fig. 1H). Finally, bulk heteroplasmy levels were positively correlated between the two systems across 164 sites shared by pairwise alignment of the zebrafish and human mitochondrial genomes, although the correlation was modest and human VAFs were systematically higher (Fig. S1K, Pearson R=0.38, P=4.2x10^-7^), indicating partially shared sequence determinants of mtDIVE activity in the two organisms.

To capture the mutational spectrum at single-molecule resolution, we performed mtFiber-seq, a long-read sequencing approach that provides full-length coverage of the 16.5 kb mitochondrial genome^36^. After 5 days of doxycycline and rapalog, individual molecules carried a median of 59 C→T mutations, showing that most mtDNA genomes had acquired multiple concurrent mutations (Fig. 1I). Parallel long-read sequencing of nuclear DNA detected no increase in C→T or G→A variant frequency after induction, placing off-target nuclear editing below the detection limit of the assay (Fig. 1J). The same was true of the nuclear genome of zmtDIVE-injected embryos (Fig. S1L). Together, these data establish mtDIVE as a flexible system for random mitochondrial mutagenesis both *in vitro* and *in vivo*.

### Metabolic context determines the mode of selection against mtDNA mutations

We next asked how metabolic context shapes the fate of mtDNA mutations and their cellular consequences. We induced hmtDIVE with doxycycline and rapalog for five days, withdrew both drugs and maintained cells in medium containing 25 mM glucose (permitting glycolysis) or 20 mM galactose (forcing reliance on OXPHOS) (Fig. 2A). Western blot analysis confirmed the rapid loss of the inducible TFAM-FKBP-DddA half upon drug withdrawal (Fig. S2A). Bulk ATAC-seq analysis revealed a marked reduction in the heteroplasmy of mtDNA variants in cells cultured in galactose (Fig. 2B), consistent with strong selection against mtDNA mutations when cells depend on OXPHOS. Heteroplasmy also decreased in glucose-rich medium, although less severely, indicating that selection operates in both metabolic contexts (Fig. 2B). To follow individual mutant molecules more precisely, we tracked two recurrent variants, G8115A (missense; *MT-CO2*) and C10205T (silent; *MT-ND4*) as representative alleles (Fig. S2B). PCR-based heteroplasmy quantification showed a significant correlation with bulk ATAC-seq (R^2^=0.73; Fig. S2C). Both variants reached ∼60% heteroplasmy after 5 days of induction. Continued growth in glucose reduced them to 30% heteroplasmy, whereas a switch to galactose rendered both nearly undetectable (Fig. S2D-E).

**Figure 2.**
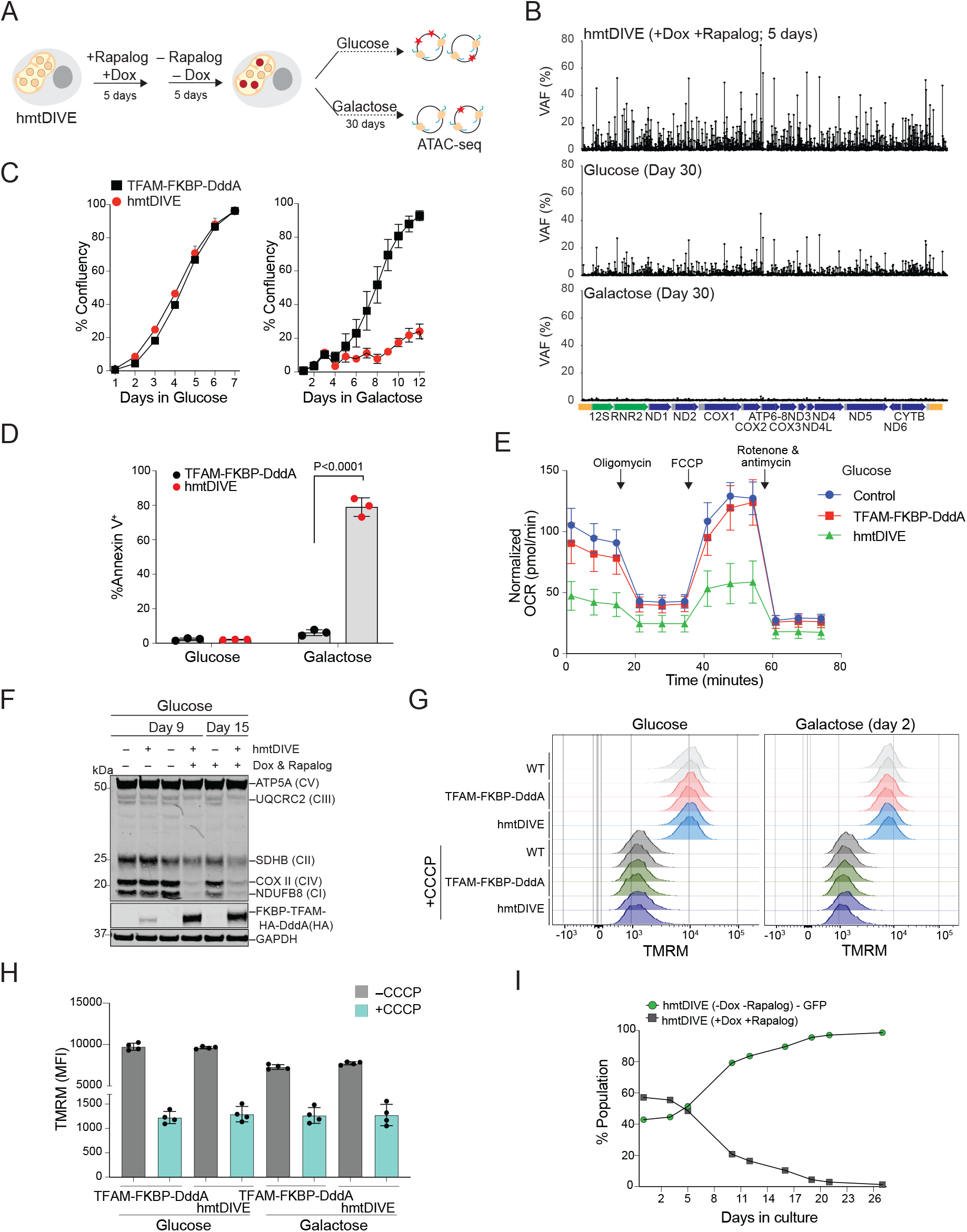
Metabolic stress and cellular competition shape the dynamics of mtDNA mutations. **(A)** Experimental design. hmtDIVE-expressing RPE cells were treated with doxycycline (Dox) and rapalog for 5 days, followed by a 5-day drug withdrawal. Cells were then maintained in glucose- or galactose-containing medium for 30 days and collected for bulk ATAC-seq analysis. **(B)** Lollipop plots showing mtDNA VAF (*y*-axis) across the mitochondrial genome (*x*-axis) in hmtDIVE cells after 5 days induction (top), and following 30 days recovery in glucose (middle) or galactose (bottom). Data are from one biological replicate. **(C)** Growth curves of cells expressing the TFAM-FKBP-DddA half (black) or hmtDIVE (red) and cultured in glucose (left) or galactose (right) after Dox and rapalog treatment. Data represent mean ± SD from *n* = 4 independent biological replicates. **(D)** Quantification of Annexin V-positive apoptotic cells cultured in glucose or galactose 2 days after induction. Cells expressing TFAM-FKBP-DddA (black) or hmtDIVE (red) were analyzed by flow cytometry. Data represent mean ± SD from *n =* 3 independent biological replicates. Unpaired two-tailed t test. **(E)** Oxygen consumption rate (OCR) measured by Seahorse in WT RPE cells (blue), TFAM-FKBP-DddA control cells (red), and hmtDIVE-expressing cells (green) cultured in glucose. Oligomycin (ATP synthase inhibitor), FCCP (uncoupler), and rotenone + antimycin A (Complex I and III inhibitors) were administered as indicated. OCR values were normalized to 30,000 cells per well. Data represent mean ± SD. *n* = 3 (WT and TFAM-FKBP-DddA), *n* = 6 (hmtDIVE) biological replicates. **(F)** Western blot analysis of representative subunits of mitochondrial respiratory chain complexes (CI-CV) in glucose-cultured cells at day 9 and day 15 after induction. HA-tagged TFAM-FKBP-DddA expression was induced with Dox (bottom panel). GAPDH was used as a loading control. **(G)** Flow cytometry analysis of mitochondrial membrane potential measured by TMRM fluorescence in WT RPE cells, TFAM-FKBP-DddA control cells and hmtDIVE-expressing cells. Cells were induced for 5 days and then cultured in glucose (left) or galactose (right) for 2 days. CCCP was used to depolarize membrane potential. DAPI staining was used to exclude non-viable cells. Biological duplicates are shown per condition. **(H)** Quantification of TMRM median fluorescence intensity (MFI) for conditions shown in (G). Data represent mean ± SD from *n* = 4 independent biological replicates. **(I)** Cellular competition assay tracking the relative abundance of induced and uninduced hmtDIVE-expressing cells over time. GFP^+^ uninduced cells (green) were mixed 1:1 with GFP^-^ cells treated with Dox and rapalog (gray) and co-cultured in glucose. The relative abundance of GFP^+^ cells was quantified by flow cytometry over time. Data represent mean ± SD from *n* = 3 biological replicates. P=0.00053, two-sided t-test of the normalized area under the change-from-baseline curve against zero.

Population-level growth analysis by IncuCyte live-cell imaging showed that mutant cells expanded comparably to controls in glucose but were markedly growth-impaired in galactose (Fig. 2C). The growth impairment was associated with extensive apoptosis, with ∼80% of cells becoming Annexin V-positive within 48 hours, whereas control cells expressing FKBP-TFAM-DddA alone remained viable in both carbon sources (Fig. 2D, S2F). Thus, forcing OXPHOS dependence triggers apoptotic elimination of cells carrying an hmtDIVE-induced mutational burden. In glucose, by contrast, mutant cells showed neither increased apoptosis nor proliferative impairment (Fig. 2C-D), yet their mtDNA mutations still declined over time (Fig. 2B), suggesting an alternative mode of selection. Seahorse assays showed that glucose-cultured hmtDIVE cells had impaired mitochondrial oxidative respiration, with reduced basal and maximal oxygen consumption alongside increased extracellular acidification rate (ECAR), indicating a compensatory shift toward glycolysis (Fig. 2E, S2G). Western blot analysis confirmed decreased expression of OXPHOS subunits spanning complexes I–IV, indicating broad disruption of the electron transport chain (Fig. 2F). These deficits were not accompanied by mitochondrial depolarization; TMRM staining showed no substantial loss of membrane potential in either glucose or galactose (Fig. 2G-H). Therefore, glucose-cultured mutant cells retained polarized mitochondria and continued to proliferate while carrying a respiratory defect, one not severe enough to trigger apoptosis. We asked whether this defect instead manifests as a competitive disadvantage, a quality-control mechanism known to eliminate metabolically impaired cells during development and in adult tissues^37^. GFP-labeled, uninduced hmtDIVE-expressing cells were cocultured at a 1:1 ratio with doxycycline- and rapalog-induced cells harboring mtDNA mutations and cultured in glucose media after drug withdrawal (Fig. 2I). Flow cytometry showed that the mutant (GFP negative) population was rapidly outcompeted, falling below <25% of by day 10 and becoming undetectable by day 26 (Fig. 2I), while control co-cultures showed no shift (Fig. S2H). CFSE CellTrace analysis independently linked mutation burden to proliferation, as cells that had undergone more divisions (low CFSE intensity) carried significantly lower G8115A heteroplasmy (Fig. S2I). These data identify two distinct selection mechanisms against mtDNA mutations, apoptosis when OXPHOS is obligatory and competitive disadvantage when glycolysis is available, which purge mtDNA mutations through different cellular processes and on different timescales.

### A mutagenic pulse triggers early respiratory compensation and a lineage-specific developmental delay

We next asked how the same mutagenic pulse plays out in a developing vertebrate, where cells differ in metabolic demand, proliferative rate and lineage identity. Survival and gross morphology were strongly dose-dependent. The highest zmtDIVE dose (50 pg) reduced survival to 24% at 48 hours post fertilization (hpf) and 15% at 72 hpf, and no surviving embryo at this dose was scored as morphologically normal at 48 hpf (Fig. 3A, S3A). Lower doses were better tolerated, with 62% and 45% of embryos injected with 20 and 35 pg appearing morphologically normal at 48 hpf (Fig. S3A). We therefore used these two doses for all subsequent analyses.

**Figure 3:**
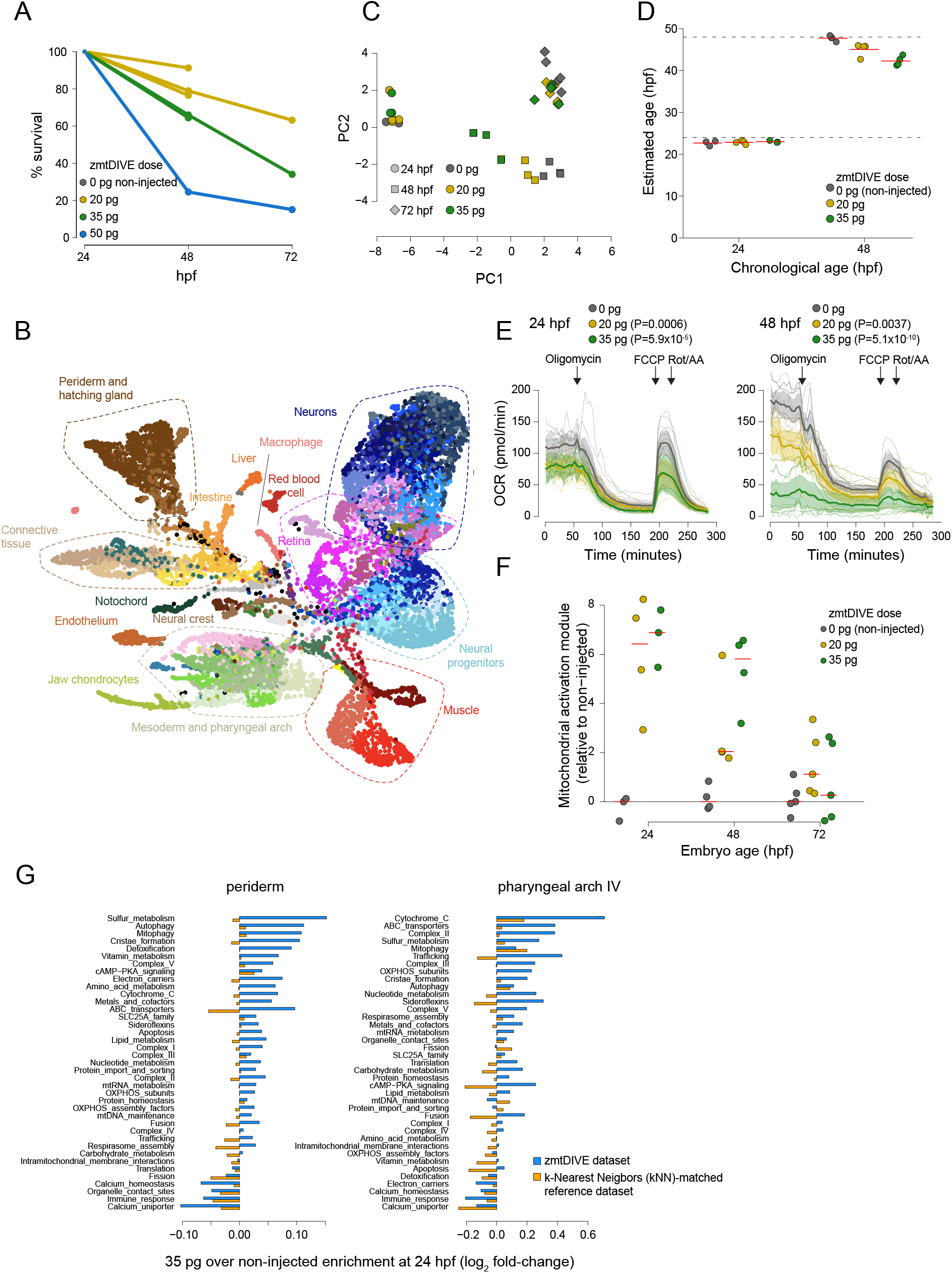
mtDIVE induces mitochondrial activation and developmental delays in zebrafish embryos. **(A)** Fraction of surviving zmtDIVE-injected embryos across RNA doses and developmental age. Each line represents an independent injection batch. N=3 (20 pg, 35 pg), 1 (50 pg) injection batches. Each batch consists of 38-72 injected embryos. **(B)** Projection of 13,010 metacells representing 551,040 single cells from 69 embryos across developmental stages and zmtDIVE doses. Colors indicate 68 annotated cell types (as in **Fig. S3D and Table S1**). **(C)** Principal Component Analysis (PCA) of pooled gene expression profiles of 37 injected embryos from Experiment 2 (see **Fig. S3E** for Experiment 1). Each symbol represents one embryo. Symbol shape indicates embryo age and color indicates zmtDIVE dose. **(D)** Estimated developmental age of 37 injected embryos from Experiment 2, based on transcriptional similarity to a reference developmental dataset^40^. Each dot represents an embryo. Dashed lines mark 24 and 48 hpf. **(E)** Oxygen consumption rate (OCR) of embryos injected with varying zmtDIVE doses, measured at 24 and 48 hpf. Arrows indicate addition of oligomycin (ATP synthase inhibitor), FCCP (uncoupler), and rotenone + antimycin A (Complex I and III inhibitors). Thin lines represent single embryos; shaded areas indicate 95% confidence intervals. n = 12 (0 pg 24 hpf), 8 (20 pg and 35 pg, 24 hpf), 12 (0, 20, 35 pg 48 hpf) embryos. P values were calculated using a linear mixed-effects model, with injection dose and inhibitor treatment as fixed effects and embryo identity as a random effect. **(F)** Aggregate expression of nuclear-encoded mitochondrial genes (derived from mitoCarta^42^) across 37 injected embryos from Experiment 2, stratified by zmtDIVE dose and developmental stage. **(G)** Enrichment of nuclear-encoded mitochondrial gene modules in 35 pg zmtDIVE-injected embryos relative to non-injected controls in the two most abundant cell types at 24 hpf (blue bars). Gold bars show the corresponding enrichment between stage-matched reference cells derived from the k-nearest neighbors (K = 10) of 35 pg–injected and non-injected embryos in the developmental reference atlas, thereby controlling for the effect of developmental delay. Values indicate log_2_ fold change.

To resolve the cellular consequences of mutagenesis we performed combinatorial-indexing single-cell RNA-seq on 428,230 cells from injected (20 and 35 pg) and control (0 pg) embryos collected at 24, 48, and 72 hpf, sampling 69 embryos across two independent experiments (Fig. 3B and Fig. S3B-C). Cells were grouped into transcriptionally homogeneous metacells^38,39^ and annotated against a published reference atlas of zebrafish development, yielding 68 cell types spanning major developmental lineages, including neurons, muscle, neural crest, and mesenchyme (Fig. 3B, S3D, Table S1, Methods)^40^. We then quantified zmtDIVE-induced changes at three levels: whole-embryo, cell-type composition, and within-cell-type (Fig. 3C, S3E-G). In parallel, we staged 24 and 48 hpf embryos *in silico* by projecting their transcriptional profiles onto a densely sampled reference dataset (Fig. 3D, Methods)^40,41^. At 24 hpf, injected embryos remained transcriptionally and compositionally similar to controls, with no discernible developmental delay (Fig. 3C-D, S3E-F). By 48 hpf, however, embryos diverged from controls in a dose-dependent manner, displaying an aggregate developmental lag of 5.7 hours at 35 pg (Fig. 3D). The delay was lineage-specific rather than uniform: mesenchymal lineages lagged by up to 8.9 hours, whereas the epidermis lagged by only 3.3 hours (Fig. S4A-B). Per-cell-type differences in developmental delays were concordant between the 20 and 35 pg doses (r = 0.82), and between independent experiments (r = 0.46, P = 0.0033; Fig. S4A-B).

The apparent developmental normalcy of 24 hpf zmtDIVE embryos masked an early respiratory perturbation. Seahorse analysis revealed a dose-dependent reduction in OXPHOS activity already at this early time point, consistent with the hmtDIVE-induced impairment observed in human cells (P=0.0006 (20 pg), 5.9x10^-5^ (35 pg), Fig. 3E). This window coincides with the period in which the zebrafish embryo derives glucose from gluconeogenesis in the extraembryonic yolk syncytial layer rather than from the liver^33^, so that embryos sustaining a respiratory deficit at 24 hpf are doing so while an endogenous, OXPHOS-independent carbon supply is active. We therefore asked whether this early metabolic perturbation was accompanied by a transcriptional footprint. Scoring nuclear-encoded mitochondrial pathways defined by MitoCarta^42^ revealed a coordinated mitochondrial activation signature in injected embryos that peaked at 24 hpf (Fig. 3F). The signature was detected in every annotated cell type and comprised induction of respiratory pathways, including cytochrome *c*, complex II and III components, cristae formation, and mitophagy, alongside suppression of calcium handling and immune-related programs (Fig. 3G, S4C). Because injected embryos are developmentally delayed, this signature could in principle reflect a shift in developmental timing rather than a direct response to mtDNA mutations. To exclude that possibility, we derived matched signatures from stage-matched cells in the reference atlas. The zmtDIVE-associated mitochondrial response was largely absent from this stage-matched reference comparison (Fig. 3G, S4C, Methods), indicating that developmental timing alone could not account for the response.

By 72 hpf the activation signature had subsided (Fig. 3F). Transcriptional divergence from controls continued to increase through 72 hpf in Experiment 1, whereas in Experiment 2 it peaked at 48 hpf and declined thereafter (Fig. 3C, S3E-F). One explanation is the selective loss of the most severely affected embryos over this window. Together, these data reveal a temporal sequence in which mtDNA mutagenesis first elicits respiratory defects and a broad mitochondrial transcriptional response while developmental progression remains largely intact, and only later, as the response subsided, produces lineage-specific developmental delay.

### Selection depletes the most heavily mutated molecules and cells

To assess how mtDNA mutation profiles vary across metabolic states and developmental stages, we jointly profiled hmtDIVE- and zmtDIVE-induced mtDNA genotypes and cellular transcription at single-cell resolution, utilizing recently published methods for mitochondria-enriched single-cell ATAC-seq with RNA-seq and sample multiplexing (DOGMA-seq, Table S2)^29,30,43^. Mutation calls showed the expected enrichment in the TC→TT and GA→AA contexts and agreed with bulk mutation frequencies (Fig. S5A). Because per-cell coverage is sparse, we developed a Bayesian framework that models per-locus editing counts as a two-component mixture and returns a posterior probability of editing for each cell and locus, thereby converting sparse counts into calibrated genotype calls (Methods).

We first applied this approach to hmtDIVE-expressing RPE cells. mtDNA mutations were induced with doxycycline and rapalog for five days, followed by three days of recovery in glucose media. Cells were then cultured in either glucose or galactose, and profiled with DOGMA-seq at days 1 and 10 (Fig. 4A). At day 1, per-cell mutation burden was comparable in both conditions. By day 10, burden had declined modestly in glucose and substantially in galactose, consistent with our findings in bulk (Fig. 4B, 2B). This decline was not distributed uniformly across the population of cells. Cells carrying the highest mutation burden (>100 detected variants) were preferentially depleted relative to medium-(10-100 variants) or low-burden (0 and 1-10 variants) cells (Fig. 4C). mtFiber-seq on cells recovered from glucose corroborated these findings, as molecules carrying 67 or more mutations were depleted relative to the induced population (Fig. 4D). Therefore, both single-cell and single-molecule measurements independently support preferential selection against cells carrying heavily edited genomes.

**Figure 4:**
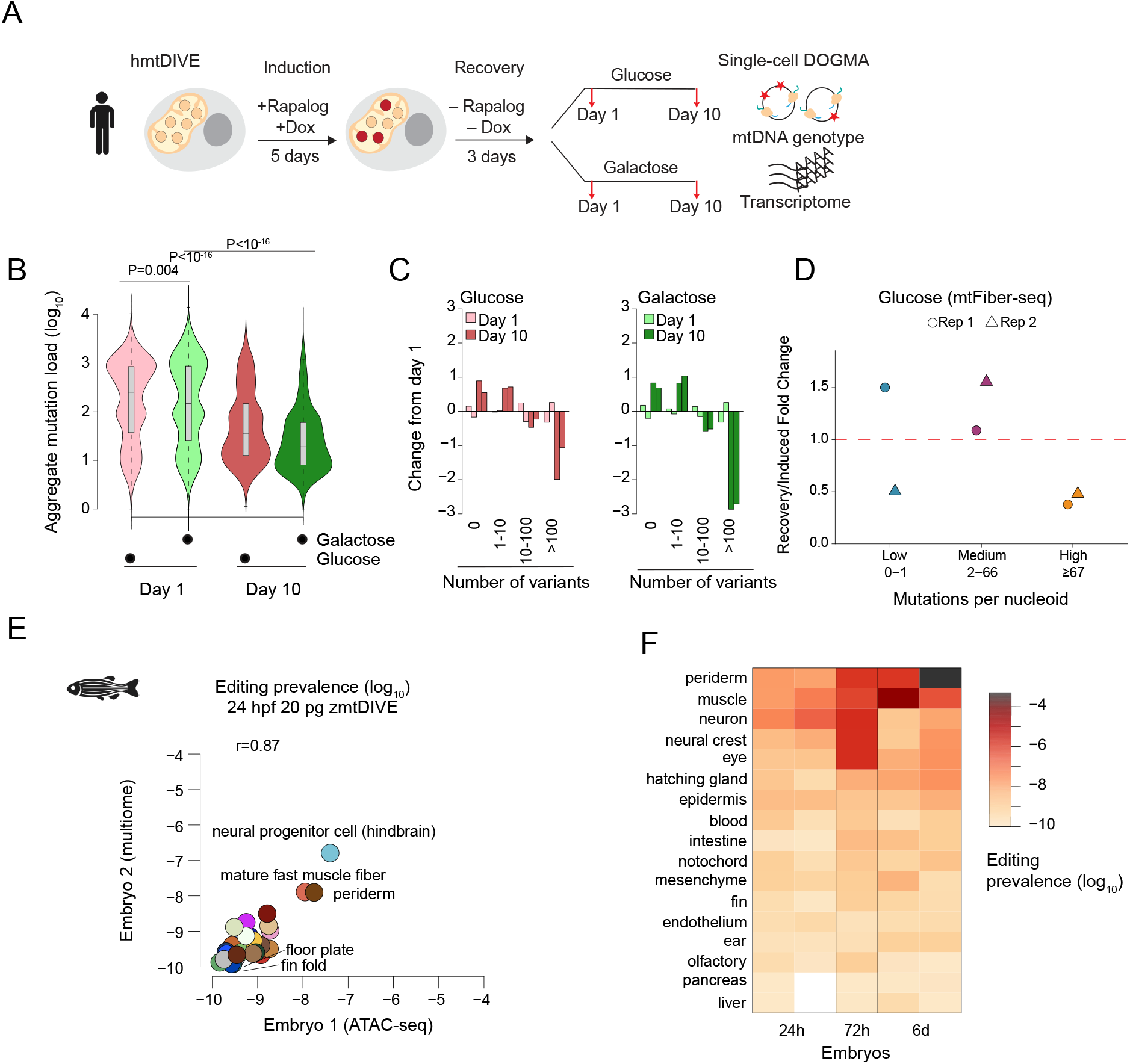
Heterogeneous mtDIVE-induced mutational load across metabolic and cell type contexts. **(A)** Experimental design. hmtDIVE-expressing RPE cells were treated with Dox and rapalog for 5 days, followed by a 3-day drug withdrawal. Cells were then cultured in glucose- or galactose-containing medium and collected for DOGMA-seq analysis after 1 and 10 days. Each condition was profiled in two biological replicates. **(B)** Distribution of aggregated mutation load across cells cultured for 1 and 10 days in glucose or galactose. Mutation load was extracted as the sum of the log Bayes factors across all TC loci in the mitochondrial genome. Two-tailed unpaired Mann-Whitney test. **(C)** Change in representation of mutation-burden categories under varying experimental conditions. Cells were stratified into none (0), low (1-10), intermediate (11-100) and high-burden (>100 variants per cell) groups. Bars represent a biological replicate; values are shown as log_2_ fold change relative to day 1. **(D)** Changes in the distribution of full-length mtFiber-seq molecules stratified by mutational burden after 10 days of culture in glucose. For each biological replicate, values represent the ratio of the proportion of recovery (day 10) molecules in each category to the proportion of induced molecules (day 1) in that category. Differences in the distribution of molecules among the three burden categories between induced and recovery samples were assessed separately for each replicate using Pearson’s chi-squared test (Rep 1: P < 2.2 × 10^-16^; Rep 2: P = 5.9 × 10^-28^). **(E)** Pearson correlation of the average prevalence of mtDNA mutations across cell types in two 20 pg zmtDIVE-injected embryos 24 hpf. The *x*- and *y*-axes show mutation prevalence in an ATAC-seq and a multiome dataset, respectively. Each point represents a cell type. **(F)** Longitudinal zmtDIVE editing activity across developmental lineages (**Table S3**), revealing lineage- and timepoint-specific differences in editing susceptibility. Columns represent 20 pg zmtDIVE-injected embryos collected from different time points. (E-F) Values represent the geometric mean fraction of cells carrying a mutation across all TC motifs.

### Cell-type-intrinsic susceptibility and lineage-specific persistence shape mtDNA editing *in vivo*

Having established that metabolic state determines which mutant cells survive, we asked whether mutation burden also differs between cell types in a developing animal. We profiled embryos injected with 20 pg zmtDIVE at 24 hpf, 72 hpf, and 6 dpf, assigning cell-type identities by reference to our transcriptional atlas (Fig. 3B, S5B-C). To compare mutation levels across samples and cell types, we down-sampled reads per cell to a fixed depth and calculated population-level mutation prevalence (i.e., the fraction of cells carrying a mutation in a given locus, Methods). Editing prevalence differed between cell types by several orders of magnitude, and these differences were reproducible across embryos (Pearson r=0.87 and 0.68 across 24 hpf and 6 dpf replicates, respectively, Fig. 4E, S5D). Cell types also differed in the ratio of mitochondrial to nuclear ATAC reads, a proxy for relative mtDNA copy number (Fig. S5E). This suggests that both mtDNA copy number and susceptibility to mtDIVE mutagenesis are properties of cell identity. However, we observed weak correlations between these two measurements, and a multivariable regression model incorporating mitochondrial read fraction, total UMI count, and total mitochondrial transcripts only partially explained the variance in editing activity across cell types, implying additional cell-intrinsic determinants of editing that these covariates do not capture (Fig. S5F-G). Notably, cell-type differences in editing prevalence affected all detectable TC editing sites similarly, pointing to a global difference in editing exposure rather than cell-type-specific site preferences (Fig. S5H-I). To follow longitudinal trends while accounting for the changing cell type repertoire during development, we grouped cells into broad developmental lineages (Table S3), and found marked differences in overall prevalence over time (Fig. 4F). Liver, olfactory, and endothelial lineages remained lowly edited throughout, whereas muscle and periderm lineages showed editing levels orders of magnitude higher. Neuronal lineages followed a distinct trajectory: editing was high at early stages but fell between 72 hpf and 6 dpf, consistent with selective loss of the most heavily edited neuronal cells over this window (Fig. 4F)^44^. Together, these analyses reveal that across both systems, mutational burden is shaped by two separable forces: metabolic demand determines how strongly heavily mutated cells are selected against, and lineage-intrinsic susceptibility governs both how frequently mutations are acquired and how long they persist during development.

### The transcriptional response to mtDNA mutations is set by metabolic and developmental context

Having established where mutations accumulate and persist, we next asked how increased mutation load reshapes the transcriptome, and whether the response depends on cellular context. To address this, we identified genes whose expression correlated with mutation load across single cells. In hmtDIVE-induced RPE cells, this depended almost entirely on carbon source. Under glucose, where energetic demand can be met glycolytically, only 22 genes passed our correlation threshold (Fig. S6A). In contrast, 825 genes (376 positively, and 449 negatively correlated) met this same criterion in galactose-grown cells (Fig. S6B). Gene Set Enrichment Analysis (GSEA) on the full ranked correlation profile also revealed differences between the two culture conditions: OXPHOS-related genes were negatively correlated with mutation load in glucose-grown cells but positively correlated when grown in galactose (Fig. S6C), consistent with a compensatory induction mounted only when OXPHOS is essential. In contrast, autophagy and mitophagy programs correlated with mutation load in both media (Fig. S6C).

A similar logic extended to development *in vivo*. In zmtDIVE-injected embryos, OXPHOS programs were positively enriched with increasing mutation load in nearly all cell types (Fig. 5A, S6D), mirroring galactose-grown human cells and suggesting that compensatory OXPHOS induction is conserved. However, beyond this shared signature, responses diverged sharply, and at times oppositely between lineages, spanning cell cycle and DNA damage pathways, phagosome and apoptosis programs, and a range of metabolic and signalling modules (Fig. 5A). Divergence was evident even within a single lineage: neuronal progenitors coupled mutational burden to DNA replication, whereas terminally differentiated neurons instead activated damage-response and apoptotic programs (Fig. 5A).

**Figure 5:**
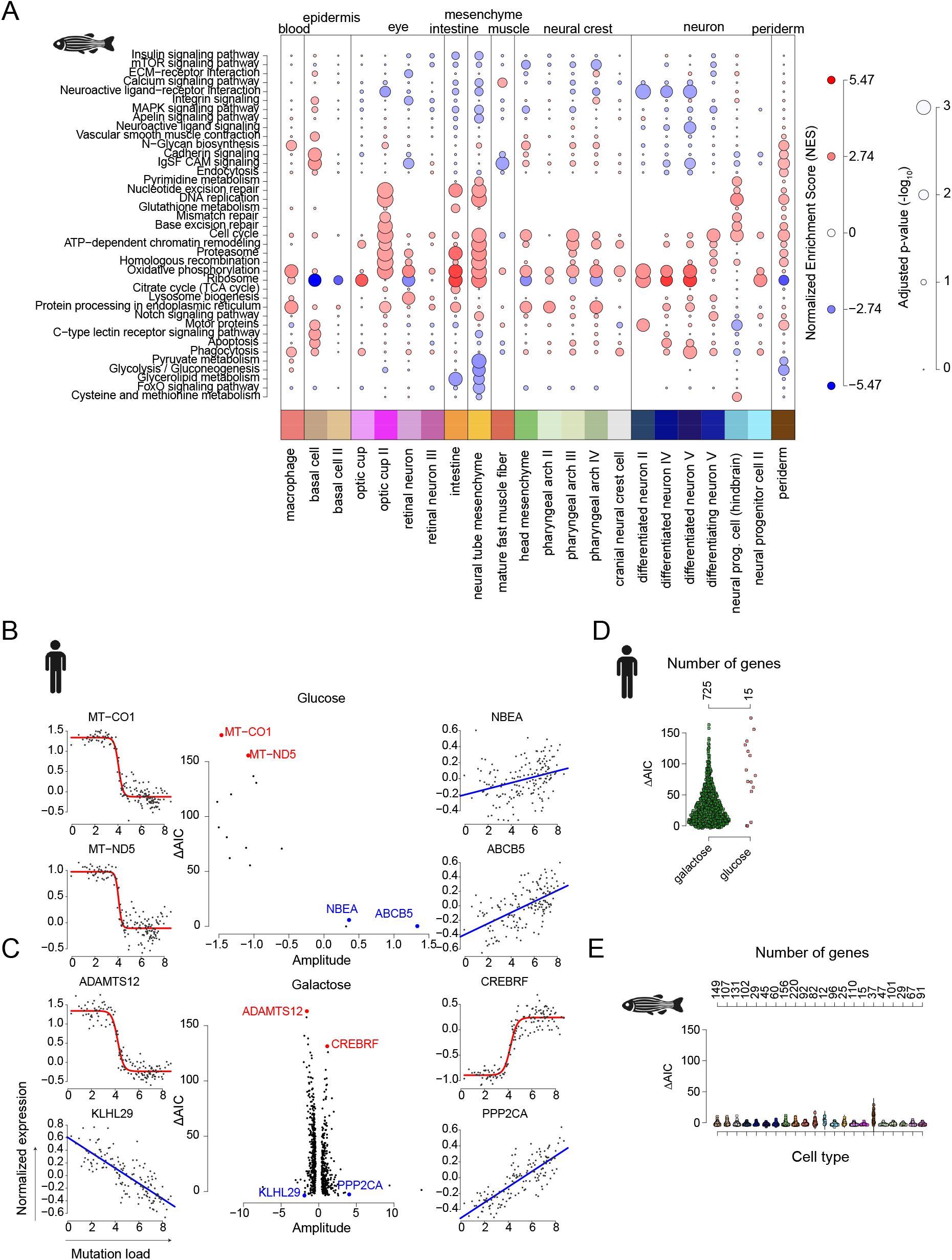
Continuous and threshold-like transcriptional responses to mtDIVE-induced mutation load. **(A)** Gene Set Enrichment Analysis (GSEA) of gene programs associated with mutation load across the 22 most abundant zebrafish cell types. Cells are pooled from 4 processed embryos. Circle color indicates normalized enrichment score (NES); circle size reflects statistical significance (FDR-adjusted p-value). **(B-C)** Classification of mutation load response genes as linear or threshold-like under glucose (B) or galactose (C). Cells were stratified by mutation load, and expression of each responsive gene (**Fig. S6A-B**) was modeled using linear and sigmoid functions of log-transformed mutation load. The *y* axis shows the difference in Akaike’s Information Criterion (ΔAIC=AIC_linear_-AIC_sigmoid_), with positive values indicating greater support for a threshold-like response curve. The *x* axis shows response amplitude. Colored dots and insets highlight genes representative at either extreme. **(D-E)** Distribution of ΔAIC values across hmtDIVE-induced cells grown in glucose or galactose (D) and across 22 most abundant zebrafish cell types (E). Each dot is a load-responsive gene. Positive ΔAIC values indicate greater support for threshold-like responses, whereas negative values indicate greater support for linear responses. Numbers of genes correlated with mutation load in each category are indicated above.

Correlation identifies which genes respond, but not how. Mitochondrial dysfunction is thought to be buffered at low heteroplasmy and to activate abruptly once a critical burden is exceeded, a behavior usually captured by a threshold or sigmoid response curve^26,31,45^. We therefore fitted each correlated gene with both a sigmoid and a linear model and compared them by ΔAIC. The Akaike information criterion (AIC) evaluates model fit while penalizing additional model complexity, allowing models with different numbers of parameters to be compared. Thus, ΔAIC indicates whether the improved fit of the sigmoid model is sufficient to justify its greater complexity relative to the linear model. In hmtDIVE-induced cells, both modes were represented (Fig. 5B-C): threshold-triggered genes, including *CREBRF* and the mitochondria-encoded transcripts, remained flat until a critical burden and then shifted abruptly, whereas burden-sensing genes such as *ABCB5* and *KLHL29* changed near-linearly across the full range. Zebrafish behaved differently; ΔAIC values were near zero in essentially every cell type (Fig. 5D-E), giving no support for a threshold model over the simpler linear one.

Two explanations could account for this difference: the smaller number of cells per zebrafish cell type, or their lower mutation load, and down-sampling would distinguish between them. Randomly sampling 1,000 human cells preserved the threshold signature, whereas restricting the analysis to the 1,000 human cells with the lowest mutation load abolished it and reproduced the flat zebrafish profile (Fig. S6E). The linear behavior in zebrafish therefore possibly reflects the lower mutational activity of mtDIVE in 20 pg embryos rather than reduced sampling depth. In summary, cells gauge mtDNA mutation load through two response modes: threshold-triggered and burden-sensing. RPE cells deploy both, whereas responses in mutagenized zebrafish scale linearly throughout, consistent with a population that remains below the threshold for abrupt activation.

### Mutational identity predicts mitochondrial transcriptional state beyond total burden

Having demonstrated that mutational burden drives context-dependent transcriptional responses, we next asked whether all mtDNA mutations contribute equally, or whether specific mutations exert distinct effects independent of overall burden. We first examined zmtDIVE-injected embryos, asking whether specific mtDNA mutations could be linked to the pan-cell-type OXPHOS program associated with high mutation burden (Fig. 5A). We trained linear regression models for each major cell lineage to quantify the contribution of individual mtDNA mutations to the emergence of this program beyond the global mutation load (Methods). Locus specificity was detected in several lineages, including eye-associated cell types, and the heavily edited muscle lineage, which exhibited the largest number of associations between individual mtDNA loci and the mitochondrial gene signature (Fig. S7A). Contributing loci were distributed throughout the mitochondrial genome, with no preference for specific mitochondrial genes (Fig. S7B). To quantify the predictive power of individual mutations, we next trained embryo-aware, cross-validated ridge regressions for each lineage, which achieved reproducible predictive performance across embryos (Fig. S7C). Although a substantial fraction of the explained variance was attributable to mutation load and technical covariates, individual mtDNA loci accounted for an additional 10-16 percentage points of explained variance (ΔR^2^, Fig. S7D) and coefficient estimates were stable across leave-one-embryo-out refits (Fig. S7E). Bootstrapped leave-one-locus-out refits showed that individual loci contributed modestly but non-redundantly, and that these contributions were lineage-specific (Fig. S7F). Global mutation load is therefore an important determinant of the transcriptional responses, but certain mutations add to that response and others attenuate it, depending on lineage context.

We next asked whether individual mutations shape the response to a defined metabolic challenge in human cells. We mutagenized RPE cells continuously for 10 days in glucose or galactose and profiled mtDNA genotypes together with nuclear transcription by DOGMA-seq. Overall, 13,847 cells passed sequencing quality control and were analyzed by combining Metacell transcriptional analysis with Bayesian inference of mtDNA mutational profiles (Methods). In contrast to the transiently induced hmtDIVE cells, highly mutated cells were recovered under both carbon sources, providing the dynamic range this analysis requires (Fig. S8A). Glucose- and galactose-grown cells occupied distinct transcriptional states (Fig. 6A), and coupled mutation load to mitochondrial transcription in opposite directions: negatively under glucose, positively under galactose, where the most heavily mutated cells sustained the highest mitochondrial expression alongside elevated stress signaling and induction of the unfolded protein response (Fig. 6B, S8B-C). Since mutation burden can in some conditions be associated with high mitochondrial transcription but with low transcription in others, we hypothesized that this distinction may reflect similarly burdened cells featuring distinct mutational profiles. We therefore tested whether mutation profiles can predict mitochondrial transcriptional state. We trained logistic classifiers to separate cells with high from low mitochondrial transcription using their full mutational profiles, fitting on one replicate and testing on the other (Fig. 6C, Methods). Full-profile models outperformed models incorporating mutation load and covariates alone (ΔAUC; Fig. 6C–D), and the gain persisted when each carbon source group was analyzed separately, strongly in galactose, more modestly in glucose, showing that distinct genotypes track divergent transcriptional states even under identical metabolic conditions (Fig. 6D). The classifier performance also transferred to independent cells collected 10 days after transient induction and to cells exposed to galactose for a single day (Fig. 6D).

**Figure 6:**
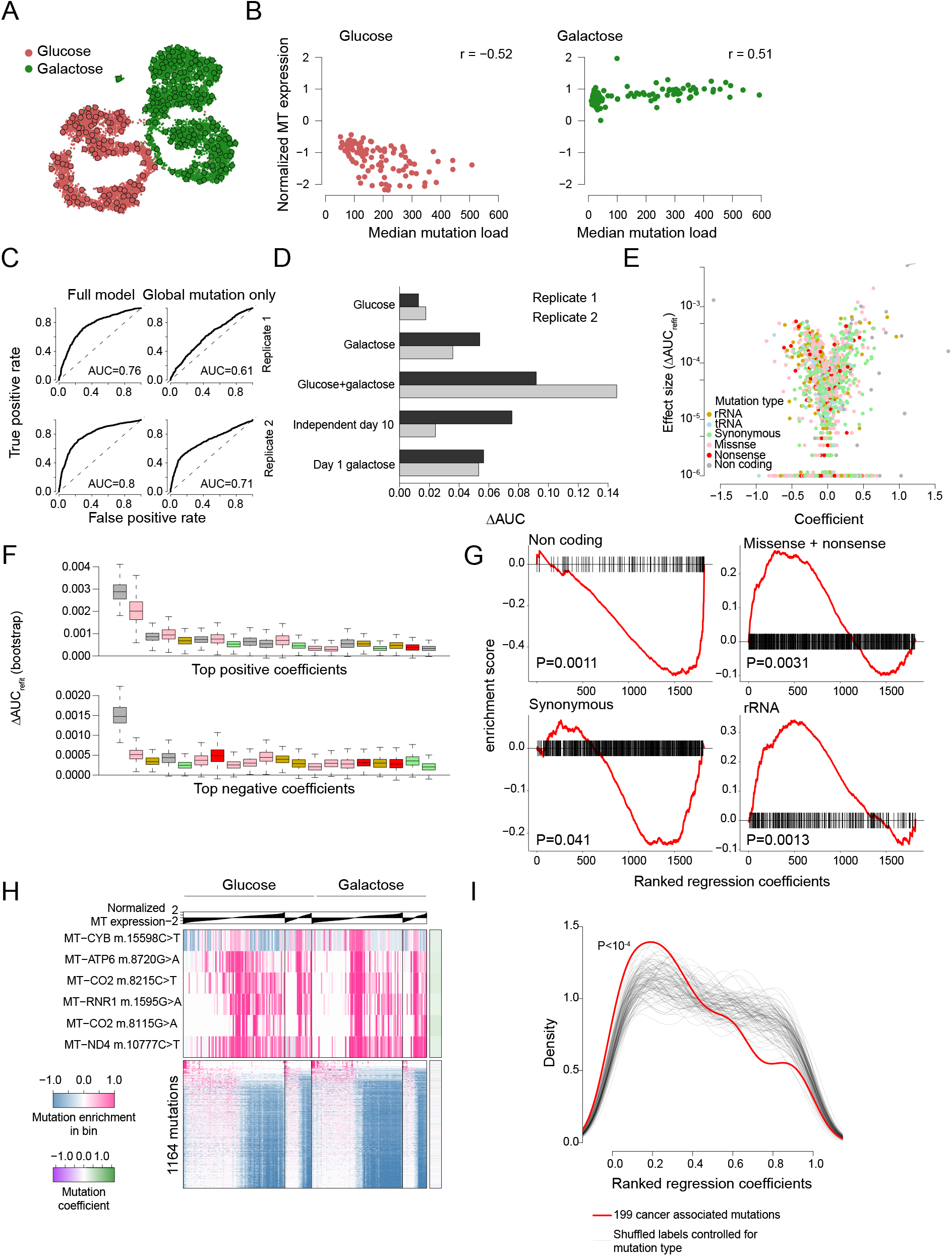
Mutational identity shapes mitochondrial transcription beyond mutation burden. **(A)** Metacell projection of 13,847 RPE cells grown for 10 days in glucose or galactose under continuous hmtDIVE induction. Each circle represents a metacell. **(B)** Pearson correlation between median mutation load (*x* axis) and mitochondrial transcription score (*y* axis) across metacells representing cells grown in glucose (left) or galactose (right). Each dot is a metacell. **(C)** Receiver Operating Characteristic (ROC) curves for logistic regression models trained on one biological replicate and evaluated on the other. Left panels show models incorporating full mutational profiles in addition to mutation load; right panels show models incorporating aggregate mutation load only. **(D)** Improvement in prediction of mitochondrial transcription state by full mutation-profile models relative to mutation-load-only models, expressed as ΔAUC=AUC_full_-AUC_load_. Bar color indicates the training replicate (1 or 2). ΔAUC was evaluated on held-out test sets comprising cells cultured in glucose, galactose or both conditions (glucose+galactose), as well as independent cohorts representing transient hmtDIVE induction after 10 days in glucose or galactose (independent day 10), and after 1 day in galactose (day 1 galactose). **(E)** Estimated effect size (*y* axis) plotted against model coefficient for individual mtDNA TC loci. Each dot represents a locus and is colored by mutation type. Effect size was computed as the drop in predictive performance (ΔAUC_refit_) following removal of the locus and model refitting. **(F)** Distribution of locus effect sizes across 500 bootstrap iterations. Shown are mtDNA loci with positive (top) and negative (bottom) model coefficients with the largest effect sizes in (E). (E-F) Colors relate to mutation location (non-coding, rRNA, tRNA, protein-coding) and type (Synonymous, missense, nonsense). **(G)** Enrichment of mutation classes along the model coefficient axis. mtDNA loci are ordered from lowest (negative) to highest (positive) coefficient. GSEA-like enrichment plots are shown for non-coding, synonymous, protein-altering (missense + nonsense) mutations and mutations in the ribosomal RNA (rRNA). **(H)** Enrichment of individual mtDNA mutations as a function of mitochondrial transcription. Day 1 hmtDIVE-induced cells cultured in glucose and galactose were binned according to mitochondrial transcription score. For each mtDNA locus and transcription bin, the 90th percentile of the log Bayes factor was calculated as a measure of mutation enrichment. Six loci with distinct enrichment patterns are highlighted (top); the remaining 1,164 variable loci are shown in the bottom panel. Rows are normalized to the median value across the first half of the transcription bins. Side color bar indicates the corresponding locus coefficient from the regression model. **(I)** Enrichment of 199 recurrent cancer-associated mtDNA mutations toward the negative pole of the model coefficient axis. mtDNA loci are ordered from lowest (negative) to highest (positive) coefficient. The distribution of recurrent mutations (red line) is compared with 10,000 permutations shuffling mutations independently within each mutation class (non-coding, synonymous, missense, nonsense, tRNA and rRNA; thin gray lines).

We next examined the contribution of individual mtDNA loci to classifier performance (Table S4). Ranking loci by the ΔAUC lost in leave-one-out refits showed that many contributed non-redundantly, with the top-ranked loci stable across bootstrap iterations (Fig. 6E-F, Methods). As in zebrafish, contributing loci were spread evenly along the genome with no enrichment for particular mitochondrial genes (Fig. S8D), but their functional classes were not. Instead, missense, nonsense and rRNA variants concentrated at the negative coefficient pole, corresponding to reduced mitochondrial transcription, and non-coding and synonymous variants at the positive pole (Fig. 6G). Strikingly, locus-specific associations with mitochondrial transcription were detected even before long-term selection had reshaped the mutational landscape. We identified six loci, all carrying positive coefficients and among those with the largest effect sizes, that were enriched in cells with high mitochondria transcription after only one day under either carbon source, in marked contrast to the remaining 1,164 variable loci, which were consistently depleted in these cells (Fig. 6H). Therefore, mutation identity is associated with distinct transcriptional states early after mutagenesis, with protein-altering variants linked to reduced mitochondrial transcription and a small subset of loci compatible with high mitochondrial output.

### Recurrent cancer-associated variants cluster at the low-transcription pole of the mtDIVE landscape

Having established that mtDNA mutation identity predicts mitochondrial transcriptional state, we next asked whether naturally occurring human mtDNA variants preferentially occupy this functional landscape. We therefore compared hmtDIVE-induced mutations with mtDNA variants identified across more than 20,000 tumors from The Cancer Genome Atlas (TCGA)^46^ spanning 33 cancer types, together with population polymorphisms from the Helix cohort^47^. Of 4,381 TCGA variants, 474 fell within hmtDIVE targetable motifs, and 97 were directly recapitulated by hmtDIVE (VAF>1%, **Fig. S9A-B**). These sites were broadly distributed across the mitochondrial genome and tumor types (**Fig. S9C**). In addition, hmtDIVE generated 547 of the 775 population polymorphisms catalogued in the Helix dataset (196,554 individuals; **Fig. S9C**), underscoring its ability to model variants found in both pathological and benign conditions^48^.

We then asked where recurrent cancer-associated mutations fall along the classifier coefficient axis defined in Figure 6^15^. Because such variants are enriched for protein-altering mutations, we tested enrichment while controlling for mutation class by permuting labels within each class. We found that the 199 mutation events recurring across multiple tumor samples and recapitulated by hmtDIVE were enriched toward the negative coefficient pole, corresponding to reduced mitochondrial transcription (P < 1x10^-4^; **Fig. 6I**). Recurrent cancer-associated mtDNA mutations therefore occupy a defined region of the functional landscape mapped by hmtDIVE, linking experimentally inferred mutation effects to patterns of mutation recurrence in human tumors^9^.

## DISCUSSION

Mitochondrial genetics has largely been practiced in reverse, whereby a candidate mtDNA variant is isolated in a cybrid or engineered in a base-edited line and examined one allele at a time^19,22–25,27^. This presumes that we already know which variants matter, and it is poorly suited to capturing the combinatorial, co-occurring mutations that real mitochondrial genomes carry. mtDIVE inverts that logic. By scattering stochastic C→T substitutions across the genome, it turns the mitochondrial genome into a substrate for pooled genetic screening, simultaneously generating thousands of variants and reading their transcriptional consequences in parallel. Here, we undertook a data-driven approach to forward genetics, anchoring on broad cellular phenotypes discernible from the transcriptome (e.g., OXPHOS program, mitochondrial transcription), and identifying individual mutations associated with them. By coupling stochastic mutagenesis to single-cell genotype-phenotype measurements, in both human cells (hmtDIVE) and an intact vertebrate (zmtDIVE), mtDIVE therefore represents a generalizable platform for forward-genetics.

Previous studies showcased the power of stochastic mitochondrial deamination in several organisms including flies and plants^49,50^. In drosophila, mitochondrially-targeted APOBEC1 generated widespread C→T substitutions and provided evidence that mutation composition can influence organismal fitness^49^. mtDIVE expands on this principle using a distinct biochemical architecture. Whereas APOBEC1 activity depends on access to single-stranded DNA, DddA modifies double-stranded DNA directly, making 5′-TC motifs throughout the mitochondrial genome accessible to mutagenesis. Fusing DddA to TFAM enables potent simultaneous mutagenesis across many such loci, generating broad combinatorial mosaicism between individual mtDNA molecules and cells. Our architecture also provides temporal and quantitative control: mtDIVE acts in a defined activity window, and heteroplasmy depends strongly on zmtDIVE injection dose or hmtDIVE induction duration. Importantly, whereas previous studies primarily resolved mutation profiles at the bulk organismal level, joint single-cell measurement of mtDNA genotype and cellular response allowed us to delineate cell-type-specific effects, compare cells carrying different mutation burdens within the same cellular context, and link individual mtDNA mutations to transcriptional phenotypes.

Mitochondrial involvement in somatic dysfunction is often framed as a burden problem, a single quantity, heteroplasmy or total load, rising until it crosses a threshold^2,4,5,7,51^. Our data challenge this view. While mutation load is useful as an indicator of overall mitochondrial dysfunction, classifiers incorporating a cell’s full mutational profile in addition to aggregate load improved prediction of the mitochondrial transcriptional state, with many loci making individually weak yet non-redundant contributions: protein-altering and non-coding variants associated with reduced and maintained mitochondrial transcription, respectively. This is the signature of a quantitative, quasi-polygenic architecture compressed into a 16.5-kb genome, akin to the shift from tumor mutational burden to driver identity. If mtDNA genotypes compositionally contribute to phenotypes, heteroplasmy is at best a coarse summary statistic, and two cells at similar aggregate loads may occupy opposite functional states.

The connection between phenotype and selection is likewise worth considering. Selection against mtDNA mutations is a property of genotype in context. Cells derived from the same mutagenic process were depleted through apoptosis when forced onto oxidative phosphorylation, but through slower competitive displacement when glycolysis provided an alternative. Moreover, 10 days of mutagenesis produced distinct genotypes across culture conditions, illustrating how metabolic demand can selectively shape mitochondrial genotypes. The zebrafish data make this apparent: individual mutations exerted phenotypic influence dependent on cell context, while mosaic mutagenesis revealed mutation dynamics consistent with differential tolerance and selection across lineages at once, such that neurons accrued high editing early then shed it, reflecting the requirement of OXPHOS in their development; while other tissues lost or maintained their mutation load^44^. The embryo’s own carbon economy may be part of what sets this balance. Zebrafish embryos generate glucose de novo in the yolk syncytial layer during precisely the stages we sampled, with glutamate as the preferred substrate and PEPCK-M as the committed enzyme^33^. A lineage’s tolerance of mtDNA damage in vivo may therefore reflect not only its intrinsic OXPHOS demand but also its access to this extraembryonic glucose supply, an in vivo counterpart to the glycolytic escape that converted apoptotic elimination into slow competitive displacement in our human cultures. This intricate interplay between mutation, selection and cell context may underlie recently described changes in mutational signatures across tissues during aging, and more broadly, shed light on why the same mutation shows such variable penetrance across patients and tissues, a puzzle heteroplasmy thresholds cannot fully explain^16,37,52^.

Perhaps the most unexpected observation is that cells carrying comparable mutational burdens respond in opposite directions. Under glucose, increased mutation load was met by reduced mitochondrial transcription and a shift toward proliferation and glycolysis. Under galactose, by contrast, heavily mutated cells instead maintained high mitochondrial transcription, arrested cell-cycle activity and engaged the unfolded-protein response. These conflicting responses suggest two coping mechanisms, whose deployment depends on whether a glycolytic escape is provided. Whether mitochondrial shutdown represents adaptive triage, limiting investment in defective machinery, or instead reflects failure to maintain mitochondrial function, is not merely semantic: the former predicts that forcing mitochondrial biogenesis would be harmful, while the latter predicts that it could be protective. This framework also offers a possible interpretation of the cancer overlap. Recurrent cancer-associated mtDNA variants were preferentially positioned at the low-mitochondrial-transcription pole. One plausible explanation is that this enrichment reflects relaxed purifying selection, rather than acquisition of adaptive traits: tumors provide a glycolytic milieu where mitochondria-damaging variants go unpurged and accumulate^9^. The same logic may extend to age-associated variants, raising the testable possibility that somatic recurrence is the record of selection switched off^53^.

## LIMITATIONS

It is important to point out that a limitation to these claims is that mtDIVE writes only at 5′-TC-3′ motifs, so its variant space is incomplete. Nevertheless, it still represents a useful model for naturally occurring and disease-associated variants, since the detected transitions are richly represented across somatic mtDNA variation, albeit with limited overlap with congenital diseases. Importantly, our current single-cell methods cannot phase co-occurring mutations onto individual mtDNA haplotypes, which constitute important units of selection and phenotypic effects. Moreover, our functional axis is transcription, a proxy of cellular activity, and the human data come from one glycolysis-tolerant RPE background that may exaggerate how permissive the glucose state is. In the future, it is possible to deploy complementary-motif deaminases to complete the variant space, defined and tunable loads to test the compositional model directly, and gain-of-function rescue to determine whether the low-expression state is triage or failure. If these hold, mtDIVE can prove instrumental in resolving what a mitochondrial genome is for a given cell, and when a cell decides it is no longer worth maintaining.

## METHODS

### Synthesis of zmtDIVE and control RNA

RNA for mtDIVE and controls was in vitro transcribed from PCR-generated templates using the mMESSAGE mMACHINE T7 Transcription Kit (Thermo Scientific). The templates of mtDIVE and controls were synthesized with Q5® High-Fidelity DNA Polymerase (New England Biolabs) by annealing DNA fragments of the 5’ (the T7 promoter) and 3’ (the poly-T tail) ends of Xenopus globin, which were PCR-generated from pT7TS (Addgene 258627), with the mt DIVE and controls sequence, respectively.

### Zebrafish handling

Zygotes of the zebrafish line TL were injected with 2 nl mtDIVE or controls (final concentration 20, 35, and 50 ng μl^-1^) (**Fig. 1B**). Zebrafish embryos were kept in E3 media (5 mM NaCl, 0.17 mM KCl, 0.33 mM CaCl_2_, and 0.4 mM MgCl_2_, pH 7.2) at 28.5°C till 5 days post-fertilization. The death and developmental defects of the embryos were quantified at 24, 48, and 72 hpf (**Fig. 3A, S3A**). Male and female zebrafish were used in each injection batch; no randomisation or blinding were done. However, dead embryos were excluded from subsequent analyses, including Seahorse and sequencing. No statistical methods were used to predetermine sample size. All animal experiments were performed in accordance with institutional and governmental regulations, approved by the Dier Experimenten Commissie of the Royal Netherlands Academy of Arts and Sciences, and conducted in accordance with the guidelines.

### Cell culture and media conditions

All human cells were maintained at 37ºC in a humidified incubator with 5% CO_2_. Human retinal pigment epithelial (RPE) cells were cultured in Dulbecco’s modified Eagle medium (DMEM, Corning, 10-013-CV) supplemented with 10% FBS, 2mM L-glutamine (Gibco), 100U/ml Penicillin-Streptomycin (Gibco), 0.1 mM MEM non-essential amino acids (Gibco). Cells stably expressing hmtDIVE were maintained in tetracycline-free medium consisting of DMEM supplemented with 10% tetracycline-free FBS (TaKaRa, 631368), 2 mM L-glutamine (Gibco), 100U/ml Penicillin-Streptomycin (Gibco), 0.1 mM MEM non-essential amino acids (Gibco), 50 µg/ml uridine (sigma-Aldrich), and 1 mM sodium pyruvate (Gibco 11360070). For experiments in galactose-containing media, cells were first seeded in tetracycline-free medium. The following day, cells were rinsed with PBS and switched to DMEM lacking glucose (Gibco, 11966025) supplemented with 10 mM Galactose (Sigma G5388), 10% dialyzed FBS (Gibco, 2640044), 2 mM L-glutamine, 100 U/mL penicillin-streptomycin,

0.1 mM MEM non-essential amino acids, 50 µg/mL uridine, and 1 mM sodium pyruvate.

### hmtDIVE molecular cloning and construct design

Plasmids encoding the split DddA cytosine deaminase halves (DddA) were fused to FKBP or FRB dimerization domains. Constructs were placed under a doxycycline-inducible TRE-tight promoter for controlled expression. For hmtDIVE, the N-terminal region of TFAM was fused in-frame to FKBP-DddA, while the complementary FRB-DddA fragment was expressed independently. Both constructs contained epitope tags (either FLAG or HA) for downstream validation by immunoblotting and immunofluorescence and a mitochondria localization signal (MLS) at the N terminus. Cloning was performed using Gibson Assembly into lentiviral backbones containing puromycin- or hygromycin-resistance cassettes. All plasmids were sequenced and verified by Sanger sequencing before use.

### Lentiviral transduction

Lentivirus was produced by transfecting 293T cells with an equal molar amount of lentiviral packing plasmids: pMDLg/pRRE (Addgene 12251), pRSV-Rev (Addgene 12253), and VSV-G (pMD2.G, Addgene 12259) in 1:1:1 ratio together with the plasmid (either TFAM-FKBP-DddA or FRB-DddA). Media was changed with fresh media on the transfected 293Ts 16 hours post-transfection. The supernatant of 293T cells containing fresh lentivirus was collected 24 hours after media change and immediately applied to cells to be infected. Infected cells are selected with the appropriate antibiotics (as described in Methods *Cell-line generation*) or sorted by flow cytometry based on the introduced fluorescent protein markers.

### Cell-line generation

Stable ARPE-19 cell lines expressing hmtDIVE were generated by lentiviral transduction. Lentivirus was produced in 293T packaging cells by co-transfection of the expression plasmid (either TFAM-FKBP-DddA or FRB-DddA) together with pMDLg/pRRE, pRSV-Rev, and VSV-G (Addgene plasmids 12251, 12253, and 12259, respectively) using a 1:1:1:1 molar ratio. The supernatant containing the virus was collected 24 hours after the media change, filtered through a 0.45 μm filter, and immediately applied to ARPE-19 cells. For controls, cells were transduced with TFAM-FKBP-DddA. Transduced cells were selected in a medium containing either puromycin (1–2 µg/mL) or hygromycin (200 µg/mL), depending on the resistance cassette encoded in the construct. Resistant colonies were expanded and maintained as polyclonal populations for subsequent experiments. Expression of hmtDIVE components was validated by Western blotting and immunofluorescence (see Methods sections *Immunoblotting* and *Immunofluorescence*).

### Induction of cells with doxycycline and rapalog

For mutagenesis induction, RPE cells stably expressing hmtDIVE or control TFAM-FKBP-DddA constructs were seeded at ∼60–70% confluency in tetracycline-free medium. The following day, cells were treated with doxycycline (1 µg/mL) to induce transcription of the TFAM-FKBP-DddA component and rapalog (50 nM; 635055, Takara Bio) to trigger dimerization of FKBP- and FRB-fused DddA halves. Unless otherwise indicated, cells were maintained under induction for 5 consecutive days with media refreshed every 48 hours to sustain drug stability. Uninduced ( – doxycycline, – rapalog) and control TFAM-FKBP-DddA cells (induced) were included in all experiments to control for potential drug-related effects.

### Immunoblotting

Cells were harvested and washed with PBS. Whole-cell extract was prepared from 1X radioimmunoprecipitation assay lysis buffer (RIPA; Millipore). Extract was clarified by centrifugation at 15,000g for 15 mins at 4°C, and protein concentration was determined by Pierce BCA (bicinchoninic acid) assay (Thermo Fisher Scientific). 8-30 µg of whole-cell extract was separated on precast 4 to 12% Bis-Tris protein gel (Invitrogen) and transferred to a nitrocellulose membrane. Membrane was blocked in PBS-TBST (137 mM NaCl, 2.7 mM KCl, 19 mM Tris Base, 0.1% Tween 20) containing 5% nonfat dry milk and incubated overnight at 4°C with primary antibody diluted in TBST-5% nonfat dry milk. Membranes were subsequently washed with TBST and incubated with appropriate IRDye 680RD and IRDye800CW secondary antibody (Li-COR Biosciences) diluted in TBST-5% nonfat dry milk. Images were detected using Odyssey Systems (LI-COR Biosciences). The following primary antibodies were used: Anti-GAPDH (CST, 2118S), Anti-Histone H3 (Abcam, ab1791), Anti-Flag (Sigma, F3165), Anti-HA (Biolegend, 901513), total OXPHOS human WB antibody cocktail (Abcam, ab110411), Anti-VDAC1 (Abcam, ab14734).

### Immunofluorescence

ARPE-19 cells stably expressing hmtDIVE were plated on coverslips in 6-well plates. Cells were stained with 125 nM MitoTracker Deep Red (Invitrogen) for 30 mins. Cells were rinsed with PBS buffer and fixed with 4% paraformaldehyde for 10 mins at RT. After PBS washes, cells were permeabilized with 0.5% Triton X-100 in PBS buffer in 10 mins at RT, followed by PBS washes. Cells on coverslips were blocked with blocking buffer (1 mg/mL BSA, 3% goat serum, 0.1% Triton X-100, 1 mM EDTA) for 30 mins at RT and incubated with anti-FLAG (Sigma, F3165) and anti-HA antibody in blocking buffer overnight at 4°C. Cells were then washed with PBS and hybridized with Alexa Fluor 488 conjugated anti-mouse or anti-rabbit secondary antibodies (Invitrogen) for 1 hour at RT in the dark. All subsequent steps were performed protected from light. After three final PBS washes, DAPI was added during the second wash to stain nuclei. Coverslips were mounted on glass slides with ProLong Gold Antifade mountant (Invitrogen). Imaging was performed using a Nikon ECLIPSE Ei2-E inverted microscope or Nikon CSU-W1 spinning disk confocal microscope.

### Subcellular fractionation

Cells were processed using CellFractionation Kit (Abcam ab109719) following standard protocol with optimization for RPE cells. 50 mL 1X Buffer A and 50 mL Buffer B (or Buffer C) were used for 500,000 cells. Buffer B was prepared by diluting Detergent I 500-fold in 1X Buffer A. Buffer C was prepared by diluting Detergent II 12.5-fold in 1x Buffer A. Cytosol and mitochondria extractions were both performed on a rotator for 30 min at RT. All centrifugation steps during the extraction were performed for 5 min at 4ºC. Cytosolic, mitochondrial, and nuclear fractions were mixed with 5X SDS-PAGE Sample Buffer and shaken at 60ºC for 10 min before western blot analysis.

### Genomic DNA purification

Total genomic DNA (gDNA) was purified by resuspending the cell pellet in 400 µL PBS containing 0.2% (w/v) SDS, 5 mM EDTA, and 0.2 mg/mL Proteinase K. The mixture was incubated at 50°C for 6 hours with constant shaking at 1000 rpm. Subsequently, DNA was precipitated by adding 0.3 M Sodium Acetate, pH 5.2, and 600 uL isopropanol and kept at -20°C for 2 hours or overnight. Precipitated DNA was centrifuged at 20,000 x g at 4°C for 30 min, followed by a 70% ethanol wash. DNA was then resuspended in TE buffer (10 mM Tris-HCl, pH 8.0, 0.1 mM EDTA) and quantified by Nanodrop.

### PCR amplification and Sanger sequencing

To validate and track specific mtDNA variants, we used primer pairs flanking two recurrent sites: G8115A (COX2) and C10205T (ND4). Primer sequences were designed to amplify ∼300–400 bp regions surrounding each variant (Forward and Reverse primers each region). PCR amplification was performed using AmpliTaq DNA polymerase (Thermo Fisher Scientific) according to the manufacturer’s instructions. Briefly, PCR reactions were carried out with 30–35 cycles under standard conditions (94 °C denaturation, 55–60°C annealing, 72 °C extension). Amplicons were purified using a PCR purification kit (e.g., Qiagen PCR Purification Kit) to remove excess primers and dNTPs. Purified products were subjected to Sanger sequencing using the forward primer for each amplicon. Sequence traces were analyzed using ICE (Inference of CRISPR Edits; Synthego) to quantify heteroplasmy levels of the target variants. As a negative control, PCR and sequencing were performed in parallel on WT RPE cells which showed no detectable mutations at the assayed loci.

### Oxygen consumption measurement (Seahorse assay)

Oxygen consumption rate (OCR) was measured using an XFe96 Extracellular Flux Analyzer (Agilent) following the manufacturer’s instructions. Cells were plated in Seahorse microplates (Agilent) at 20,000 cells per well 24 hours before the measurement. After the cells adhered to the plate, cell culture media were removed and replaced with Seahorse media (DMEM, Agilent 103575) supplemented with 10 mM glucose and 4 mM glutamine. OCR analysis was performed at basal conditions and after oligomycin (1 µM), FCCP (1 µM), or rotenone plus antimycin A (each at 1 µM) treatment, as indicated, according to the manufacturer’s instructions.

### Oxygen consumption measurement of live zebrafish embryos

Seahorse analysis of live zebrafish embryos was performed following an established protocol^54^. The Seahorse cartridges (102,416, Agilent) were hydrated by adding 1000 μl of XF calibrant solution to the utility plate and incubated overnight at 28.5°C. At 24 hpf or 48 hpf, the chorion was removed using thin forceps. Single embryos at 24 hpf and 48 hpf were then transferred using a plastic Pasteur pipette to each well of the Seahorse XF 24 Islet Capture Microplate (101122-100, Agilent) in 525 μl E3 medium at 28.5°C. The baseline of embryonic respiration was measured in 12 cycles, with each cycle consisting of 1 min mixing, 1 min waiting, and 2 min measuring. After measuring basal respiration, the drugs oligomycin (9.4 µM, 30 cycles), FCCP (8 µM, 6 cycles), and rotenone/antimycin A (1.5 µM each, 15 cycles) were sequentially added to each well. After the assay, the genomic DNA of each embryo was extracted using the DNeasy kit (69504, Qiagen), and its relative mtDNA copy number was measured by qPCR with primers for the mtDNA MT-ND1 and with primers for the nuclear B2-microglobulin, as previously described^55^. The normalization of the experiments is based on the relative mtDNA copy number obtained.

### Growth analysis by Incucyte

Cells were seeded in Tet-free medium into 24- or 48-well plates at 7,000 to 10,000 cells per well, with three wells per experimental condition. After 12 hours (when cells were fully attached), cells were rinsed with PBS and replaced with treatment media (glucose or galactose medium; DMEM containing 25 mM glucose or galactose, supplemented with 10% dialyzed FBS, 4 mM glutamine, 50 µg/mL uridine, 1% PenStrep, 1 mM sodium pyruvate). Plates were imaged every 12 hours on the Incucyte S3 Live-cell analysis system using brightfield and a 10x objective. Cell confluency was measured using Incucyte analysis software.

### Cell competition assays

GFP-labeled and unlabeled hmtDIVE cells were used to perform a competition assay. GFP^+^ hmtDIVE cells that had not been exposed to doxycycline/rapalog (“untreated controls”) were mixed at a 1:1 ratio with GFP^-^ hmtDIVE cells that had undergone 5 days of doxycycline and rapalog treatment to induce mutagenesis. Cells were seeded at equal density and maintained under standard culture conditions in glucose-containing media. Mixed cultures were passaged every 3-4 days at ∼80% confluency to avoid density-dependent artifacts. At each passage (days 0, 5, 10,15, 20, and 27), cells were collected and analyzed by flow cytometry. Control co-cultures were established by mixing GFP^+^ and GFP^−^ untreated hmtDIVE cells at a 1:1 ratio to rule out any growth differences caused by fluorescent labeling.

### CFSE CellTrace proliferation assays

Cell proliferation was assessed using CellTrace CFSE dye (Thermo Fisher Scientific, C34554), which stably labels cytoplasmic proteins and is diluted by half with each cell division. hmtDIVE cells were first treated with doxycycline (1 µg/mL) and rapalog (50 nM) for 5 days to induce mtDNA mutagenesis. Following induction, cells were harvested, resuspended in PBS containing 0.1% BSA, and stained with CFSE according to the manufacturer’s protocol (5 µM final concentration, 20 min at 37 °C). Staining was quenched by adding five volumes of complete medium and incubating for 5 min at 37 °C before washing.

CFSE-labeled cells were seeded in glucose-containing media and cultured for 7 days. At the end of the assay, cells were harvested and analyzed by flow cytometry (BD LSRFortessa). CFSE fluorescence intensity was measured in the FITC channel, and data were analyzed with FlowJo software to determine the distribution of dye dilution across cell generations. To link proliferation history with mitochondrial mutation status, cells were stratified by CFSE intensity, and heteroplasmy for the G8115A variant was quantified by targeted PCR and Sanger sequencing (see *PCR amplification and Sanger sequencing*).

### Mitochondrial membrane potential and apoptosis analysis

Cells cultured in Tet-free medium were stained with 1 mM Tetramethylrhodamine, Methyl Ester, Perchlorate (TMRM, Thermo Fisher Scientific T668) in culture media for 30 min at 37ºC. Cells were then harvested by trypsinization, centrifuged, resuspended in 3% FBS in PBS, and analyzed on an LSR Fortessa analyzer. TMRM signal was detected with a 561 nm laser and a 586/15 Bandpass filter. Recorded data were analyzed using FlowJo software. As a positive control, WT cells were treated with 20 mM CCCP for 6 hours. For apoptosis analysis by Annexin V staining, cells were seeded in equal numbers in a Tet-free medium. The next day, cells were rinsed with PBS and changed to glucose- or galactose-medium. We harvested cells 24-48 hours after media change. A culture medium containing floating dead cells was collected with trypsinized cells. Cells were stained with the Annexin V (Pacific Blue) detection kit and 7-AAD (BioLegend 640926) per the manufacturer’s instructions. Cells were analyzed on LSR Fortessa analyzer. Recorded data were analyzed using FlowJo software.

### Bulk deep-profiling of single embryo mutational profiles

Single zebrafish embryos were treated with proteinase K, followed by a nested PCR strategy to amplify regions of mtDNA (17 amplicons, **Table S5**). For library preparation, a PCR reaction was performed that introduced only Illumina TruSeq adapters. All libraries were sequenced paired-end at 75 bp read-length on the Illumina NextSeq platform.

Raw reads from multiplexed PCR-based deep sequencing assays were processed as follows: Adapter sequences (AGGACGACGGCAACTACAAG) were trimmed from read 1 using cutadapt (v2.6). Reads were aligned to the zebrafish mitochondrial genome (GRCz11) using bwa (v0.7.18). Resulting alignments were sorted and indexed using samtools (v1.21), and per-base coverage was computed. Pileups were generated using bcftools (v1.21).

Variant Call Format (VCF) files were converted to tabular format using a custom script. Amplicon intervals were defined based on primer sequences, and positions overlapping primer-binding sites were excluded from downstream analysis to avoid PCR-induced artifacts. mtDNA loci with coverage below 100× were excluded.

Substitution frequencies were calculated for each locus as the fraction of reads supporting a non-reference base out of total coverage. When aggregating substitution frequencies across loci, geometric means were used to account for the wide dynamic range of editing levels. Geometric means were computed with a small pseudocount added to avoid undefined values; the pseudocount magnitude was adapted to the expected expression range in each analysis.

Estimates of editing levels in the nuclear genomes were derived from single-cell mt-enriched ATAC-seq data (see below, section *Processing of mt-enriched scATAC-seq, multiome, and DOGMA-seq data*).

### Bulk ATAC-seq of human RPE cells

50,000 cells were collected and frozen in 10% DMSO in culture media in -80°C before submission to the Integrated Genomics Operation (IGO) core at Memorial Sloan Kettering Cancer Center. Library preparation followed standard ATAC-seq protocol without nuclei isolation^56^.

### Mitochondrial variant calling – Bulk ATAC-seq

We called mitochondrial DNA variants in the bulk-ATACseq data as previously described in Kim et al^57^. To determine high-confidence heteroplasmic mitochondrial genotypes, we employed a computational variant-calling pipeline that incorporates individual per-base, per-allele base quality (BQ) scores. Variants were identified using two complementary approaches: Mutect2 (GATK v4.1.2.0, mitochondrial mode) and samtools mpileup (v1.9). Only variants detected by both pipelines were retained for downstream analysis. For samtools mpileup, we required a minimum mapping quality (MAPQ > 20) and base quality (BQ > 20), with at least 2 alternative-supporting reads observed on both the forward and reverse strands. PCR duplicates and low-quality reads were removed prior to variant calling. Variants falling within blacklisted homopolymer regions of the mitochondrial genome (positions 513-525 and 3105-3109) were excluded. To minimize false positives, strand concordance was assessed, and variants were required to show Pearson correlation coefficient ≥ 0.2 between forward and reverse reads. This approach enabled accurate detection of heteroplasmic variants, and we verified the reproducibility of our calls by comparing variant profiles across multiple sequencing runs. Final calls represent high confidence heteroplasmic genotypes suitable for downstream analyses.

For **Fig. 1H**, editing coverage was quantified as the fraction of DddA-editable mtDNA sites (5′-TC-3′ and reverse-complement 5′-AG-3′ motifs) at which a corresponding C>T or G>A variant was detected at VAF ≥ 1%. Fractions were calculated independently for each biological replicate and are shown as mean ± SD across three independent replicates.

### Comparison of hmtDIVE and zmtDIVE editing patterns

Human and zebrafish mtDNA positions were matched through pairwise alignment of mitochondrial protein-coding sequences at the peptide level. For each protein-coding gene, human and fish peptide sequences were extracted and aligned using pairwiseAlignment (Biostrings v2.54.0) with the BLOSUM62 substitution matrix. Aligned peptide positions were mapped back to their corresponding nucleotide coordinates, resulting in a total of 164 homologous loci with a conserved TC motif across species. Substitution frequencies at these loci were compared between a human bulk ATAC-seq sample and a pseudo-bulk scATAC-seq sample from a 20 pg zmtDIVE-injected zebrafish embryo at 24 hpf (**Fig. S1K**).

### Fiber-seq long-read sequencing

Mitochondrial isolations and mtFiber-seq were performed as previously described^36^. ARPE-19 cells were grown in 150 mm plates to 80–90% confluence. Cells were pelleted by spinning at 150 × g for 5 minutes at 4°C. Media was removed, and cells were resuspended in 1 mL Cell Lysis Buffer (10 mM Tris, pH 7.5; 10 mM NaCl; 1.5 mM MgCl_2_). Cells were allowed to swell on ice for 7.5 minutes, then lysed by douncing in a 7 mL glass dounce with 20 strokes. A 2 M Sucrose T10E20 Buffer (10 mM Tris, pH 7.6; 1 mM EDTA, pH 8.0; 2 M Sucrose) was added to bring the final sucrose concentration to 250 mM.

Cell debris was pelleted by centrifuging lysates at 1,300 × g for 3 minutes at 4°C. The supernatant was transferred to fresh tubes and spun again at 1,300 × g for 3 minutes at 4°C. The supernatant was then transferred to fresh tubes, and mitochondria were pelleted by spinning at 18,000 × g for 15 minutes at 4°C. Mitochondrial pellets were resuspended in Permeabilization Buffer (20 mM Tris, pH 7.4; 70 mM Potassium Acetate; 250 mM Sucrose; 0.25% Tween-20; 0.25% NP-40/Igepal/CA-630) and incubated on ice for 10 minutes. Mitochondria were then pelleted by centrifugation at 18,000 × g for 15 minutes at 4°C.

Supernatant was removed, and pellets were resuspended in 56 µL mtFiber-seq Reaction Buffer (20 mM Tris, pH 7.4; 70 mM Potassium Acetate; 250 mM Sucrose). Then, 1.5 µL S-adenosylmethionine was added to a final concentration of 0.8 mM, and tubes were transferred to a thermocycler prewarmed to 37°C. Reactions were allowed to proceed for 10 minutes at 37°C before quenching with 3 µL 20% SDS. Sample volume was increased to 200 µL by adding 7 µL 20% SDS and buffer. Protein was degraded by adding 2 µL of 18.2 mg/mL Proteinase K (Millipore Sigma 3115887001) and incubating at 55°C for 1 hour.

Phenol:chloroform:isoamyl alcohol extraction was performed to extract DNA, followed by ethanol precipitation using 0.1 volumes Sodium Acetate and 2.5 volumes ethanol. DNA was pelleted, washed with 70% ethanol, and air dried for 10 minutes at room temperature before resuspending in 81 µL water. Then, 10 µL NEB CutSmart Buffer and 3 µL RNase A (Thermo Fisher Scientific AM2270) were added along with restriction enzymes. For all samples, 3 µL XmaI (NEB R0180S) was added to fragment chromatin, and an additional 3 µL BamHI-HF (NEB R3136L) was added to linearize mtDNA.

Reactions were incubated at 37°C for 1 hour. A phenol:chloroform:isoamyl alcohol extraction was performed, followed by chloroform:isoamyl alcohol extraction to remove residual phenol. DNA was precipitated as before, pelleted, washed with 70% ethanol, and air dried. DNA was quantified using Qubit hsDNA (Thermo Fisher Scientific Q32851) and sequenced on a PacBio Sequel IIe system at the University of Maryland Institute for Genome Sciences.

### PacBio Single-Nucleotide Variant Calling

Single-nucleotide Variants (SNVs) for individual PacBio reads were identified using a custom pipeline. CIGAR strings of pbmm2 aligned CCS BAM files were parsed to extract the position within the read, the appropriate reference genome position, and the mismatched base within the read. The reference genome position was used to extract the reference base from chrM_BamHI.fa. Data were written to a .tsv indicating the individual read name, the number of mismatches, the read and reference positions of each mismatch, and the read and reference identities of each mismatched position. Mutation positions were converted to standard rCRS coordinates. Positions less than or equal to 2,307 were adjusted by increasing by 14,262. Otherwise, the position was decreased by 2,307. Cell-type-specific SNVs were determined as those present in >95% of reads within the uninduced sample and were removed from all samples.

### PacBio Off-target Nuclear Deamination Analysis

Potential off-target DddA-induced cytosine deamination in nuclear DNA was assessed by analyzing CT variant calls using DeepVariant (v.1.2.1) output VCFs generated against the GRCh38/hg38 reference genome. All CT transitions and their corresponding allele frequencies were extracted from each sample. Allele frequencies were compared between treated and control samples using the Wilcoxon rank-sum test to detect enrichment of CT variants indicative of off-target nuclear deamination events. High-frequency variants (AF > 50%) representing cell-line single-nucleotide variants were retained in the analysis to preserve the complete variant spectrum. Statistical significance was set at p < 0.05.

### Combinatorial-indexing scRNA-seq (ScaleBio)

Zebrafish embryos at 24 hpf, 48 hpf, and 72 hpf were dissociated into single cells by 0.25% trypsin, filtered through 40-μm cell strainers, and resuspended in DPBS. Cells were then fixed with ScaleBio (now 10X Genomics) ScalePlex Fixation Kit according to the vendor’s instructions. Each embryo was associated with a unique ScalePlex oligo for sample demultiplexing. In case the cell amount from dissociated 24 hpf embryos did not reach the minimum required cell input for the fixation kit (min. 50,000 cells per fixation reaction), multiple embryos were pooled into one fixation reaction. After fixation, individually ScalePlexed samples were pooled into a single sample pool and stored at -80°C until further processing. The scRNA-seq library construction and sequencing were performed by Single Cell Discoveries B.V. using the ScaleBio Single Cell HT kit v1.1 with the ScalePlex library add-on, according to the vendor’s instructions. Libraries were sequenced on an Illumina NovaSeq X Plus instrument, targeting 20,000 reads per cell for scRNA libraries and 2,000 reads per cell for ScalePlex libraries, with an estimated capture of 500,000 cells.

### Processing of combinatorial indexing single-cell RNA-seq data

Low-level processing of combinatorial indexing scRNA-seq data was performed using ScaleRna (v2.0.0). Reads were demultiplexed by sample and library structure, followed by adapter trimming based on protocol-specific combinatorial barcodes and linker sequences defined for the ScalePlex and Quantum library chemistries. Trimmed reads were aligned to the zebrafish reference genome (GRCz11) using STAR.

Cell-containing barcodes were identified using the cellFinder method (FDR < 0.001), yielding 581,340 candidate cells. Cells with fewer than 500 or more than 20,000 total UMIs were excluded, resulting in 566,013 cells.

Assignment of cells to individual embryos was performed based on ScalePlex barcode recovery matrix. Cells were retained only if the two most abundant ScalePlex barcodes matched expected row and column identifiers, yielding 551,040 cells for downstream analysis. Retrieved cells had a median of 2,412 UMI and 1,394 genes per cell (**Fig. S3C**).

### Cell type annotation using reference atlas mapping

Reference expression matrices were obtained from a densely sampled scRNA-seq atlas of early zebrafish development^40^. As this dataset includes Perturb-seq experiments, only cells from non-perturbed embryos were retained. For each developmental stage in the query dataset (24, 48, and 72 hpf), reference cells from the matching stage were selected.

Expression matrices were imported using monocle3 (v1.4.26). Label transfer from the reference to the query dataset was performed using Seurat (v5.3.0), applying the FindTransferAnchors and TransferData functions with the first 30 principal components. Reference annotations, including tissue, broad cell type, and detailed cell type labels (metadata fields cell_type_broad and cell_type_sub), were transferred to query cells.

Transferred annotations were subsequently refined through metacell-based clustering (see below).

### Metacell construction and transcriptional clustering

Metacell clustering was performed using the Metacells Python package^39^. An iterative clustering strategy was applied, in which metacell partitions were generated, evaluated, and refined across successive rounds. Default parameters were used unless otherwise specified.

To reduce the influence of non-cell-type-specific transcriptional programs, we first defined a set of anchor genes associated with general cellular processes, including cell cycle (*mcm3, mcm4, mcm7, mki67, pcna, ube2c, top2a, cdk1*) and immediate-early response (*fos, jun, egr1, hsp90ab1, txn, nr4a3*). The mark_lateral_genes function was used to identify additional genes exhibiting strong correlation with these anchors, representing candidate lateral gene modules. These genes were subsequently curated and excluded from downstream clustering.

Metacells were constructed using the divide_and_conquer_pipeline and collect_metacells functions. Following an initial round of clustering, additional noisy and lateral genes were manually identified and excluded (*col2a1a, col11a1a, col1a2, co1a1a, col1a1b, postnb, slc4a4b, myhz1*.*1, znf185, znf185, and1, and2, muc5*.*1, ENSDARG00000076830, TTN, smyhc1, ENSDARG00000058556, ENSDARG00000069559, ENSDARG00000105341, ebf3a, and ptprua*). The dataset was then re-clustered using the filtered gene set. Metacells contained a median of 39 cells (12-96). 8,333 outlier cells identified by the Metacells pipeline were discarded from downstream analysis.

Each metacell was assigned an initial annotation based on the most frequent transferred cell type labels (broad and subtypes) of its constituent cells. To refine these annotations, we used published marker genes for each cell type^40^ and computed their enrichment across metacells (**Table S1**). Marker enrichment profiles were manually inspected to validate and, where necessary, adjust metacell annotations.

### Analysis of transcriptional changes following zmtDIVE injection

Global transcriptional changes following zmtDIVE injection were quantified using three complementary approaches:

1. *mbryo-level pseudo-bulk analysis*. For each embryo, gene expression profiles were obtained by pooling all cells and computing the geometric mean expression per gene. Log_2_ fold changes per gene were calculated relative to the median expression across embryos. The top 25 most enriched genes per embryo were selected, and the resulting embryo-by-gene matrix was subjected to principal component analysis (PCA) (**Fig. 3C, S3E**).
2. *Cell type composition analysis*. For each embryo, cell type proportions were computed and normalized. Log_2_ fold changes of each cell type were calculated relative to the median across embryos, and the resulting embryo-by–cell type matrix was analyzed using PCA (**Fig. S3F**).
3. *Cell type-specific transcriptional perturbations*. For each cell type, cells were grouped by embryo, and gene expression profiles were computed as in (1). Only embryo and cell type pairs with at least 50 cells were retained. Log_2_ fold changes relative to the median across embryos were calculated, and genes showing at least two-fold enrichment were considered. The top 25 enriched genes per embryo were used to construct a feature matrix for PCA. Euclidean distances in the resulting PC space (first five components) were computed, and for each embryo, the median distance from non-injected controls was reported (**Fig. S3G**).

### In silico developmental staging of embryos

Embryo developmental age was inferred in a similar fashion to Dorrity et al.^41^, by mapping query cells onto a densely sampled reference scRNA-seq atlas of zebrafish development, in which embryos were profiled at 2-hour intervals^40^. For each tissue-level annotation, reference and query cells sharing the same label were subset and analyzed independently. Only reference cells with annotated ages up to 72 hpf were considered.

Mapping of query cells to the reference was performed using Seurat (v5.3.0) functions FindTransferAnchors and MapQuery. For each query cell, the k nearest neighbors (k = 10) were identified in the reference dataset using FNN (v1.1.4.1; get.knnx), based on the first 30 principal components of the reference.

Query embryos at 72 hpf were excluded from downstream analysis due to reduced sampling density in the reference beyond 48 hpf. For each query cell, a transcriptional age was defined as the mean annotated age of its 10 nearest reference neighbors. Cell-type-specific developmental ages were then computed by averaging transcriptional ages across cells of the same type within each embryo.

To obtain a cell-type-aware estimate of developmental delay, baseline ages were defined for each cell type within each condition (developmental stage and experimental batch) using non-injected controls. For each cell, the deviation from its corresponding baseline age was calculated, and these values were aggregated across all cells within each embryo to derive an overall estimate of developmental delay (**Fig. 3D**).

### Gene module enrichment analysis of mitochondrial pathways

Mitochondrial gene annotations were obtained from MitoCarta3.0 (MitoPathways3.0)^42^. Human genes were mapped to zebrafish orthologs using biomaRt (v2.42.0). In total, 1,002 genes assigned to 40 functional groups were identified in the scRNA-seq dataset. Of these, 966 genes with at least 1,000 total detected UMIs were retained for downstream analysis.

A mitochondrial activation score (“mito-score”) was computed for each cell as the geometric mean expression of the retained genes. To obtain cell-type-aware estimates of mitochondrial activation, baseline mito-scores were defined for each cell type within each condition (developmental stage and experimental batch) using non-injected controls. For each cell, deviations from the corresponding baseline were calculated and aggregated across all cells within each embryo to derive an embryo-level mito-score (**Fig. 3F**).

To assess pathway activation independently of developmental delays, we leveraged the kNN-based mapping to the developmental atlas described above. For each cell type and embryo, observed gene expression was defined as the geometric mean expression across all cells. To derive a developmentally matched reference, all reference cells corresponding to the k-nearest neighbors of query cells were pooled (with repetition), and their geometric mean expression was computed (“controlled expression”). Pathway-level expression was then calculated by averaging gene expression within each mitochondrial functional group, and compared between observed and controlled expression profiles (**Fig. 3G, S4C**).

### mtDNA-enriched single-cell ATAC and Multiome profiling

#### Human cells: modified DOGMA-seq

Single Cell Multiome ATAC + Gene Expression of human cells was performed with the 10X Genomics system using the Chromium Next GEM Single Cell Multiome Reagent Kit A (catalog no. 1000282) and the ATAC Kit A (catalog no. 1000280). We followed the Chromium Next GEM Single Cell Multiome ATAC + Gene Expression Reagent Kits User Guide, with the exception that the single-cell suspension was prepared as described in Mimitou et al. (DOGMA-seq LLL protocol with NP-40 buffer). However, to further increase mitochondrial DNA coverage, we optimized the lysis step by increasing NP-40 concentration to 0.3% and the time of lysis to 3 min. Final libraries were sequenced on the Illumina NovaSeq X Plus platform (R1-28 cycles, i7-8 cycles, R2-90 cycles).

#### Zebrafish embryos: mt-enriched scATAC-seq and Multiome

Zebrafish embryos at 24 hpf, 72 hpf, and 6 dpf were dissociated into single cells by 0.25% trypsin, filtered through 40-μm cell strainers, and resuspended in DPBS. Samples for mt-enriched 10x ATAC-seq and multiome kits were then processed as previously described^58^. In brief, cells were fixed with 1% formaldehyde for 10 min at room temperature, which was then quenched with 0.125 M glycine. Cells were washed and centrifuged twice in 1 ml PBS at 400g for 5 min at 4°C. Chilled lysis buffer with 0.3% NP-40 (100 µl) was then added to the loosened cell pellet, and the mixture was gently pipetted up and down three times. The cells were incubated on ice for 3 min, washed with the wash buffer, and then centrifuged at 500g for 5 min at 4 °C. After discarding the supernatant, the pellets were resuspended in chilled 1× diluted nuclei buffer, and cell concentrations were counted.

### Processing of mt-enriched scATAC-seq, multiome, and DOGMA-seq data

Low-level processing of mt-enriched single-cell datasets was performed using cellranger-atac (v2.1.0) for scATAC-seq samples and cellranger-arc (v2.0.2) for multiome and DOGMA-seq samples. In both cases, reads were aligned to a custom reference genome based on the zebrafish GRCz11 build, or to the human reference genome downloaded from the cellranger website. Fragment files were aggregated across libraries using the aggr function to ensure uniform processing.

Following alignment and barcode demultiplexing, BAM files were separated into mitochondrial and nuclear reads. Per-cell read counts were computed for each compartment to derive mitochondrial capture efficiency metrics. Mitochondrial BAM files were further split into single-cell BAMs using the list of high-confidence cell barcodes provided by Cell Ranger. Cells with less than 4,000 mitochondria-aligned reads were discarded from subsequent analysis.

Single-cell-aware pileups were generated using bcftools mpileup (v1.21) with the read-group option. Resulting BCF files were converted into tabular format using custom scripts. Given the low background error rates and the downstream application of a Bayesian mixture model to distinguish signal from noise, no additional base-quality or strand-bias filtering was applied at this stage. Coverage tracks were computed using samtools depth. When required, reads were downsampled to a fixed depth (4,000 reads per cell) using the samtools subsample option.

In the DOGMA-seq dataset, cells were assigned to biological samples based on the HTO barcode with the highest UMI count. Only cells with >50 UMIs assigned to the most abundant HTO and a >2-fold ratio between the most abundant and second-most abundant HTO UMI counts were retained for downstream analysis.

To compare mtDNA and nuclear editing rates in zmtDIVE-injected embryos, a large nuclear genomic region (Chr3:13,979,574–41,938,720) was selected, and bulk-level pileups were generated using bcftools (**Fig. S1L**).

### Cell type annotation of zebrafish multiome datasets

Multiome datasets of zebrafish embryos were first mapped to the combinatorial indexing reference dataset (**Fig. 3**) using Seurat (v4.1.1). The 24 hpf sample was mapped to 24 hpf reference cells, while 72 hpf and 6 dpf samples were mapped to 72 hpf reference cells. Label transfer was performed using FindTransferAnchors and TransferData with the first 30 principal components.

scATAC-seq datasets were subsequently mapped to the annotated multiome cells. Peak matrices from both modalities were aggregated, and joint dimensionality reduction was performed using Signac (v1.2.1) with RunTFIDF and RunSVD. For each ATAC cell, the k nearest neighbors (k = 10) in latent semantic indexing (LSI) space were identified from the multiome dataset, and cell type identities were assigned by majority voting.

### Bayesian modeling of mtDNA editing rates across TC motifs

We applied Bayesian inference to derive estimates for editing levels across mtDNA loci, while controlling for variable coverage and sequencing errors. We modeled per-locus editing counts as a two-component mixture of negative binomial distributions. Negative binomial distributions were preferred over binomial distributions to account for overdispersion in editing counts across loci and cells, which deviated substantially from binomial expectations. For each cell i and locus j:

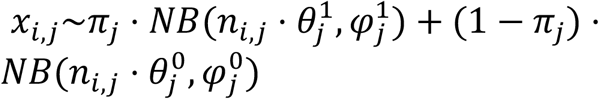

Where X_i,j_ is the number of reads supporting a substitution in j, n_i,j_ is the total coverage in j, θ^1^ and φ^1^ denote the mean editing rate and dispersion in the edited component, θ^0^ and φ^0^ the background mean and dispersion, and π represents the mixing proportion. We placed weakly informative priors on all parameters, using uniform Beta(1,1) priors for the editing rates (θ^1^, θ^0^) and mixing proportion (π), and Gamma(2,0.1) priors for the dispersion parameters (φ^1^, φ^0^).

We used rstan (v2.32.6) to infer posterior distributions of editing rates and mixing proportions for each TC motif separately in each embryo. n_i,j_ coverage was derived from per-cell coverage tracks (using samtools depth), and X_i,j_ substitution counts from pileup data. Initial value assignments for θ^0^ were informed from the non-injected zmtDIVE control ATAC-seq sample. In hmtDIVE cells, we used a constant 0.001 initial assignment for θ^0^. Inference was performed in 4 chains with 5000 iterations per chain. When θ^0^ exceeded θ^1^, component labels were switched such that the higher-θ component represented the edited state, and the corresponding mixture proportion was changed from π to 1−π.

Posterior distributions were used to derive binary editing matrices. For each posterior draw, we computed the likelihood of the data under the edited and background components (P_1_ and P_0_ respectively), and defined the posterior probability of editing *P* = *P*_1_/(*P*_0_ + *P*_1_). A cell i was considered edited at locus j if P>0.95 in the majority of posterior samples and the number of edited reads was greater than 1. The posterior threshold (P > 0.95) was selected to ensure high-confidence editing calls, while the majority rule across posterior samples increased robustness to sampling variability. The resulting binary matrices were used to quantify the total number of variants per cell (**Fig. 4C**), and to derive empirical editing prevalence across cell types and embryos (**Fig. 4E-F, S5D**).

In addition, we computed the log Bayes factor (lBayes)=log_10_(P_1_/P_0_) as a continuous measure of confidence in editing events relating to each cell and locus.

Global mutation load per cell (as in **Fig. 4B, 6B**, etc.) was defined as the sum of positive lBayes values across the 1,000 (zebrafish) or 1,800 (human) most frequently detected mtDNA loci. This threshold was chosen empirically, as mutation load saturated beyond it, robustly separating edited from non-edited cells.

Results were insensitive to the exact number of loci used.

### Association of gene programs with mtDNA mutation load

To identify gene programs associated with mtDNA mutation load in hmtDIVE-induced cells under glucose and galactose, we stratified them into 30-cell bins based on their aggregate mutation load. For each gene and bin, we calculated the geometric mean of the gene’s expression in the bin. Gene-wise Pearson correlations between expression and median mutation load were computed across bins.

To identify gene programs associated with mtDNA mutation load across zebrafish cell types, we restricted analysis to cell types represented by at least 400 cells. Within each cell type, cells were partitioned into transcriptionally homogeneous metacells using the Metacell R package (v0.3.7). Gene features were selected using variance-to-mean and coverage-based filters (mcell_gset_filter_varmean and mcell_gset_filter_cov; parameters: T_vm=0.01, T_tot=5, T_top3=3), and metacells were constructed using mcell_mc_from_coclust_balanced (K=20, min_mc_size=20). Metacell-level mutation load was computed as the median across constituent cells.

Gene expression values were summarized at the metacell level and transformed to log_2_ fold-change relative to the median expression within each cell type. Gene-wise correlations between expression and mutation load were computed across metacells. To ensure robust signals, we retained genes with ≥ 50 total UMIs and sufficient dynamic range (third-highest enrichment >0.15 within the cell type).

Gene Set Enrichment Analysis (GSEA) was performed on ranked correlation profiles using zebrafish KEGG pathways (retrieved via KEGGREST v1.26.0, org.Dr.eg.db v3.10.0 for zebrafish and org.Hs.eg.db v3.10.0 for human cells), implemented with fgsea (v1.12.0, 10,000 permutations).

### Distinguishing between linear and threshold-like gene activation

To classify transcriptional responses to mutation load as continuous or threshold-like, we first stratified cells into bins according to aggregate mutation load. Bins contained 30 cells for human data and 15 cells for the less abundant zebrafish cell types. For each cell gene and mutation-load bin, we calculated the geometric mean expression (with pseudocounts of 0.1) and median mutation load. Responsive genes were defined as those with expression variance >0.05 across bins and absolute Pearson correlation >0.3 between expression and mutation load (**Fig. S6A–B**). For zebrafish, this analysis was restricted to cell types containing at least 400 cells. The number of responsive genes identified in each zebrafish cell type is shown in **Fig. 5E** (top axis).

For each responsive gene, expression across mutation load bins was fitted using both a linear model and a four-parameter sigmoid function: 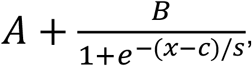, where A represents the baseline expression, B the response amplitude, c the mutation-load threshold and s the scaling factor controlling response steepness. Initial parameter values were set to A=min(y), B=max(y)-min(y), c=median(x), s=0.5*standard deviation(x), where x denotes mutation load and y denotes gene expression. The Akaike Information Criterion (AIC) of both models was calculated and the ΔAIC=AIC(linear)-AIC(sigmoid) metric was used to determine the better fit, such that positive values indicate greater support for a threshold-like response.

### Linear modeling of locus-specific contributions controlling for global mutation burden across zebrafish lineages

For each zebrafish lineage, we tested whether individual mtDNA loci were associated with transcriptional responses beyond global mutation load. Analyses were restricted to heavily edited cells, defined as cells with aggregate mutation load ≥ 5. An oxidative phosphorylation gene module was defined using the corresponding KEGG pathway, and per-cell module scores were calculated using Seurat’s AddModuleScore function, followed by centering and scaling within each embryo (OXPHOS score).

We retained mtDNA loci with detectable editing in at least 0.1% of cells. For each locus, lBayes values were divided by the 99th percentile across cells, and truncated to the [0,1] interval, thereby reducing the influence of extreme values. To control for global mutation load while excluding the contribution of the focal locus, a self-excluded mutation load was calculated for each cell by summing editing values across all loci except the locus being tested.

For each locus, we fitted a linear model of the form:

OXPHOS score ∼ locus + excluded_load + embryo + mtDNA_read_count + mtDNA/nuclear read ratio where the focal predictor is the normalized lBayes value of the locus, with embryo identity included as a categorical covariate. Regression coefficients and FDR-adjusted p-values were extracted for each locus (Fig. S7A-B).

### Regularized ridge regression for multi-locus genotype–phenotype prediction

To evaluate the joint contribution of mtDNA loci to the OXPHOS score, we fitted a separate ridge regression model for muscle and eye lineages using glmnet (v4.1-10) with the following design: OXPHOS score ∼ 1,732 loci + mutation_load + mtDNA_read_count + mtDNA/nuclear read ratio, where for each locus and cell, its observed editing value was defined as (1 + number of mutated reads) / (1 + coverage). Values for each locus were normalized to the maximum across analyzed cells. For each lineage, cells were first partitioned into 80% training and 20% held-out test sets, with proportional representation from each embryo. Within the training set, embryo identity was used to define cross-validation folds for selection of the ridge penalty parameter. Predictive performance was evaluated on the held-out test set. To quantify the predictive information contributed by locus identity beyond aggregate mutation burden and technical covariates, we trained a covariate-only model (excluding loci) in parallel and computed (on the held-out test set): ΔR^2^=R^2^(full model) - R^2^(covariates only) (**Fig. S7D**). Robustness of model coefficients across embryos was assessed by explicit leave-one-embryo-out refitting, and coefficient consistency across refits was evaluated (**Fig. S7E**). To estimate the locus-specific contributions to prediction, we performed bootstrapped leave-one-locus-out refitting. In each bootstrap iteration, the full model was fitted and its test-set R^2^ calculated. Each locus was then removed individually, the model was refitted, and the locus-specific decrease in predictive performance was calculated as: ΔR^2^(refit)=R^2^(full)−R^2^(−locus).The distribution of ΔR^2^(refit) values across 500 bootstrap iterations, representing the magnitude and robustness of locus-specific contributions, is shown in **Fig. S7F** for the eye and muscle lineages.

### Building classifiers predicting mitochondrial transcriptional states from individual mutations

hmtDIVE cells were classified into mitochondrial transcription states (MT^low^ and MT^high^). A mitochondrial transcription score was calculated from the aggregate expression of the 13 mtDNA-encoded protein-coding genes and standardized across the complete scRNA-seq dataset, comprising continuous and transiently induced hmtDIVE cells collected at days 1 and 10, thereby defining MT-low and MT-high states on a common transcriptional scale across induction regimes, time points, and culture conditions. Cells with negative or positive z-scores were classified as MT^low^ and MT^high^, respectively. Consequently, representation of the two states was not necessarily balanced within individual samples.

We trained two ridge-penalized logistic regression classifiers using glmnet, corresponding to reciprocal training and testing between the two day-10 continuous hmtDIVE replicates including cells from glucose and galactose culture conditions. Models had the following design:

MT_state ∼ 1814 loci + mutation_load + mtDNA_read_count + mtDNA/nuclear read ratio

Analyses were restricted to cells with aggregate mutation load >0, after z-score scaling across all cells. For each locus, its observed editing value was defined as (1 + number of mutated reads) / (1 + coverage). Values for each locus were normalized to the maximum across analyzed cells. For each classifier, the ridge penalty parameter was selected by five-fold cross-validation within the training replicate, with fold identity assigned randomly. Classification performance on the held-out replicate was quantified as area under the receiver operating characteristic curve (AUC) using the ROCR package (v1.0-11). To quantify predictive information contributed by locus identity beyond aggregate mutation load and technical covariates, we fitted a covariate-only model (excluding loci) in parallel and calculated: ΔAUC=AUC(full model) - AUC(covariates only) (**Fig. 6C-D**).

To quantify locus-specific contributions to classifier performance, each locus was removed individually and the classifier refitted. Locus-specific effect size was calculated as:ΔAUC(refit)=AUC(full)−AUC(−locus). ΔAUC_refit was calculated once for all loci using the held-out test replicate (**Fig. 6E**). Uncertainty was subsequently assessed by 500 bootstrap iterations for the 20 positive-coefficient and 20 negative-coefficient loci with the largest effect sizes (**Fig. 6F**). The leave-one-locus-out and bootstrap analyses were performed using the classifier trained on replicate 1 and evaluated on replicate 2.

Coefficients from the two fitted classifiers were averaged for each locus to define the functional axis of mtDNA variation. Enrichment analysis of mutation classes along this ranked axis was performed using fgsea (v1.12.0) with 10,000 permutations (**Fig. 6G**). Enrichment of 199 recurrent cancer-associated mutations along the same axis was compared with a null distribution generated from 10,000 permutations in which mutation labels were shuffled independently within each mutation class, thereby preserving mutation-class composition (**Fig. 6I**).

### Cross-dataset comparison of hmtDIVE-induced variants with cancer and population databases

To assess the disease and population relevance of hmtDIVE-induced mutations, we compared high-confidence variants generated in hmtDIVE cells with variants cataloged in The Cancer Genome Atlas (TCGA) and the HelixMTdb dataset.

hmtDIVE-induced variants were defined as sites with variant allele frequency (VAF) ≥ 1% in bulk ATAC-seq after 5–14 days of induction. Variants were aggregated across replicates and only positions consistently detected in at least two experiments were retained for downstream analyses. TCGA variants (n = 4,381) were obtained from published mitochondrial variant calls across >20,000 tumor samples spanning 33 cancer types. Variants were filtered to include only canonical C>T and G>A transitions, which represent the most frequent cancer-associated mtDNA substitutions. Helix variants (n = 775) were obtained from HelixMTdb, a population database of mtDNA variation from 196,554 individuals. For each site, we computed the total number of carriers (TOTAL_COUNT = HET_COUNT + HOM_COUNT) and the proportion reported as heteroplasmic (FRACTION_HET = HET_COUNT/TOTAL_COUNT). These values represent the fraction of individuals in the cohort carrying a given variant in heteroplasmic form versus homoplasmic form. Importantly, Helix does not provide per-individual heteroplasmy levels (i.e., percent mutant load within each carrier), only whether a carrier is categorized as heteroplasmic or homoplasmic. For consistency, we restricted comparisons to variants with population allele frequency ≥ 0.01%.

Control region (D-loop) variants were excluded from both TCGA and Helix analyses because this region contains multiple hypervariable sites with elevated background mutation rates and no reproducible phenotypic associations. Overlap was determined by direct coordinate matching against the revised Cambridge Reference Sequence (rCRS, NC_012920).

### Quantification and statistical analysis

Statistical analysis of the experiments was performed using GraphPad Prism or R. Sample sizes, replicate numbers, and statistical tests are reported in the corresponding figure legends. Values are reported as mean ± SD as indicated in the figure and supplementary figure legends.

## Data availability

Raw and processed sequencing data used in this study have been deposited on GEO and will be made available as of the date of publication under accession numbers: GSE346506, GSE346508, GSE347326, and GSE349286.

## Code availability

The code used to process the data and generate all figures was archived in Zenodo and will be made available as of the date of publication.

## Declaration of generative AI and AI-assisted technologies in the manuscript preparation process

During the preparation of this work, the authors used ChatGPT and Claude for edits and proofreading. After using these tools, the authors reviewed and edited the content as needed and they take full responsibility for the content of the published article.

## Acknowledgments

We thank members of the Sfeir and van Oudenaarden labs for commenting on the manuscript. We thank Toby Lieber for experimental assistance, and Xilin Jiang and Amos Tanay for advice. We thank Single Cell Discoveries for their help with single-cell sequencing services.

## Author Contributions

Conceptualization: AG, RB, CYC, AS, AVO

Methodology: AG, CYC, JP, RB, RuC, ED, MEK, RoC, RSI

Investigation: AG, CYC, RB, MK, JA, RaS, ML, RoS, JP, RC, RSI

Funding acquisition: DP, ER, CBT, RSI, AVO, AS

Project administration: AVO and AS

Supervision: AVO and AS

Writing – original draft: AG and AS

Writing – review & editing: all authors

## Competing interest

Agnel Sfeir is a co-founder, consultant, and shareholder of Repare Therapeutics.

This communication represents the views of a member of the PERSIST-SEQ consortium and neither the consortium as a whole nor the European Union and EFPIA are liable for any use that may be made of the information contained herein.

## SUPPLEMENTARY FIGURE LEGENDS

**Figure S1:**
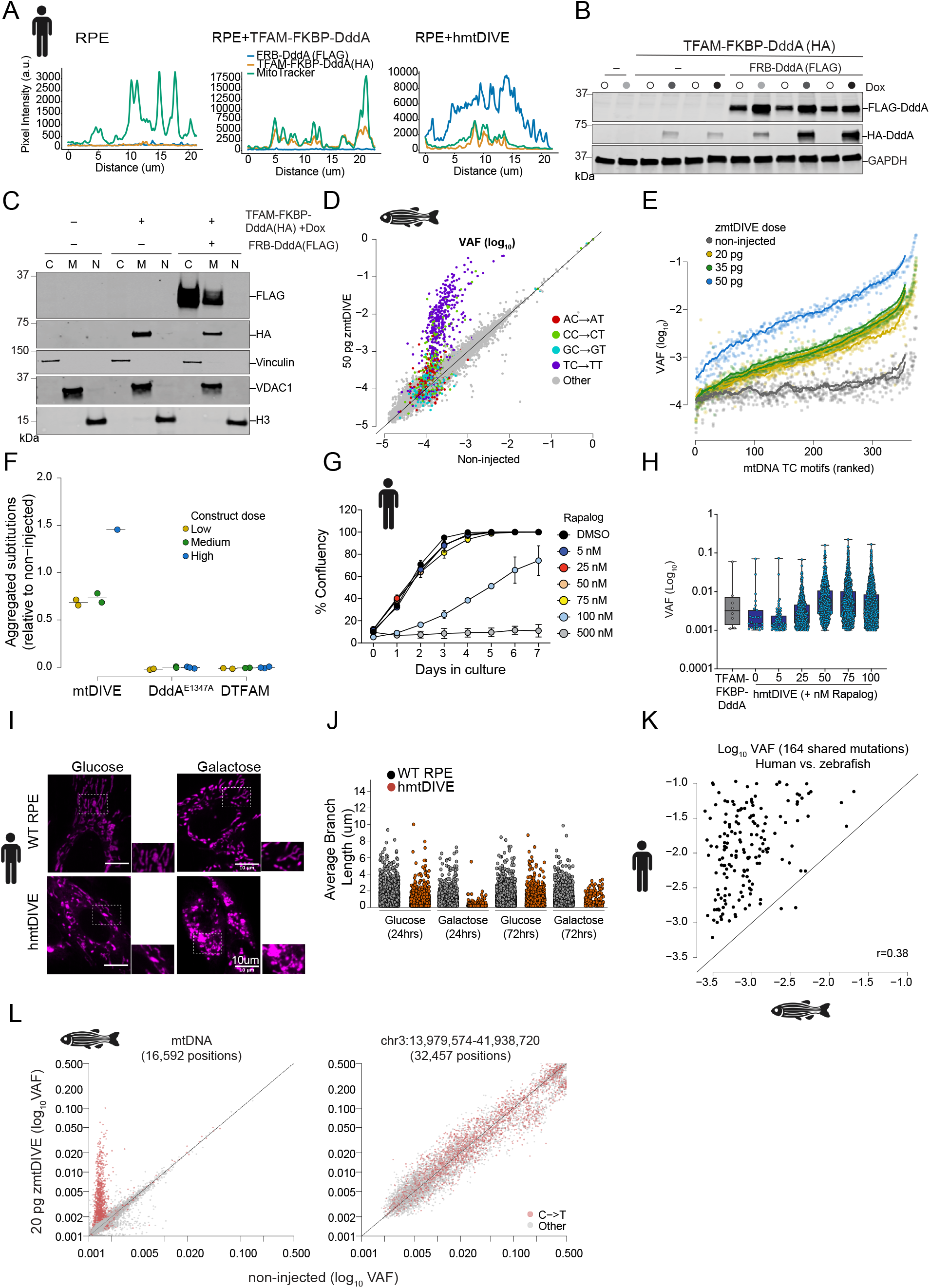
Benchmarking mtDIVE across organisms. **(A)** Line scan analysis of hmtDIVE component localization. Pixel intensities were measured along the yellow lines indicated in Fig. 1D for WT RPE, TFAM-FKBP-DddA, and hmtDIVE-expressing cells. Profiles show signal intensities for FRB-DddA (FLAG-tagged, blue), TFAM-FKBP-DddA (HA-tagged, orange), and Mitotracker (green) along the indicated axis. Scale bars, 10 µm. **(B)** Western blot validating hmtDIVE expression in ARPE-19 cells under increasing concentrations of doxycycline. HA- and FLAG-tagged fusion proteins were detected; GAPDH was used as a loading control. **(C)** Subcellular fractionation and immunoblotting confirming mitochondrial enrichment of hmtDIVE components. Cytoplasmic (C), nuclear (N), and mitochondrial (M) fractions were probed for HA and FLAG. Vinculin, H3, and VDAC1 served as compartment-specific markers. **(D)** Comparison of substitution frequencies (log_10_ VAF) across all detected mtDNA positions between non-injected embryos (0 pg; *x*-axis) and embryos injected with a high dose of zmtDIVE (50 pg; *y*-axis) 24 hours post fertilization (hpf). Colored dots represent C→T substitutions, with colors indicating the upstream sequence context. **(E)** Substitution frequencies (log_10_ VAF) across all detected TC motifs in embryos injected with varying zmtDIVE doses at 24 hpf. Each line represents a rolling-window average for one embryo; motifs are sorted by their mean substitution frequency across all injected embryos. **(F)** Mean zmtDIVE activity in 24 hpf embryos injected with mtDIVE, with zmtDIVE with an inactive DddA variant and without the TFAM domain. Each point is an embryo. Values represent the log_10_ geometric mean of substitution frequencies across all detected TC motifs relative to non-injected controls. **(G)** Cell confluency dynamics measured by IncuCyte live-cell imaging over 7 days in cells treated with increasing concentrations of rapalog (DMSO, 5-500 nM). Data represent median ± SD from independent biological replicates (*n* = 3). **(H)** Distribution of mtDNA variant allele frequencies (VAFs) in hmtDIVE-expressing cells treated with increasing concentrations of rapalog (1–100 nM) for 5 days. Treatment with 500 nM resulted in complete cell death and was not analyzed. Variants are plotted as log_10_(VAF); each dot represents a mtDNA variant, averaged across *n* = 3 replicates. TFAM-FKBP-DddA-expressing cells are shown as controls. Box plots show the median (center line), interquartile range (box), and whiskers extending to 1.5x the interquartile range. **(I)** Live-cell images of mitochondrial networks stained with MitoTracker in WT RPE, TFAM-FKBP-DddA, and hmtDIVE-expressing cells cultured in glucose or galactose. Scale bars, 10 μm. **(J)** Quantification of mitochondrial branch length in WT RPE, TFAM-FKBP-DddA, and hmtDIVE-expressing cells under the indicated metabolic conditions (glucose or galactose, 24 hours and 72 hours). Measurements were performed using the MiNA ImageJ macro on live MitoTracker-stained cells. A total of *n =* 60 cells per condition. Data represent one biological replicate per condition. **(K)** Pearson correlation between bulk substitution frequencies (log_10_) of shared TC motifs in hmtDIVE-expressing RPE cells (y axis) and zebrafish embryos injected with 20 pg zmtDIVE (*x* axis). Shared identified shared mtDNA loci by amino-acid pairwise alignment of mitochondrial protein-coding sequences. **(L)** Comparison of substitution frequencies across detected mtDNA positions between non-injected embryos (0 pg; *x*-axis) and embryos injected with 20 pg zmtDIVE (*y*-axis), assessed by bulk ATAC-seq. Shown are 16,592 positions across mtDNA (left), and 32,457 positions within a nuclear region on chromosome 3 (right). Colored dots represent C→T substitutions.

**Figure S2:**
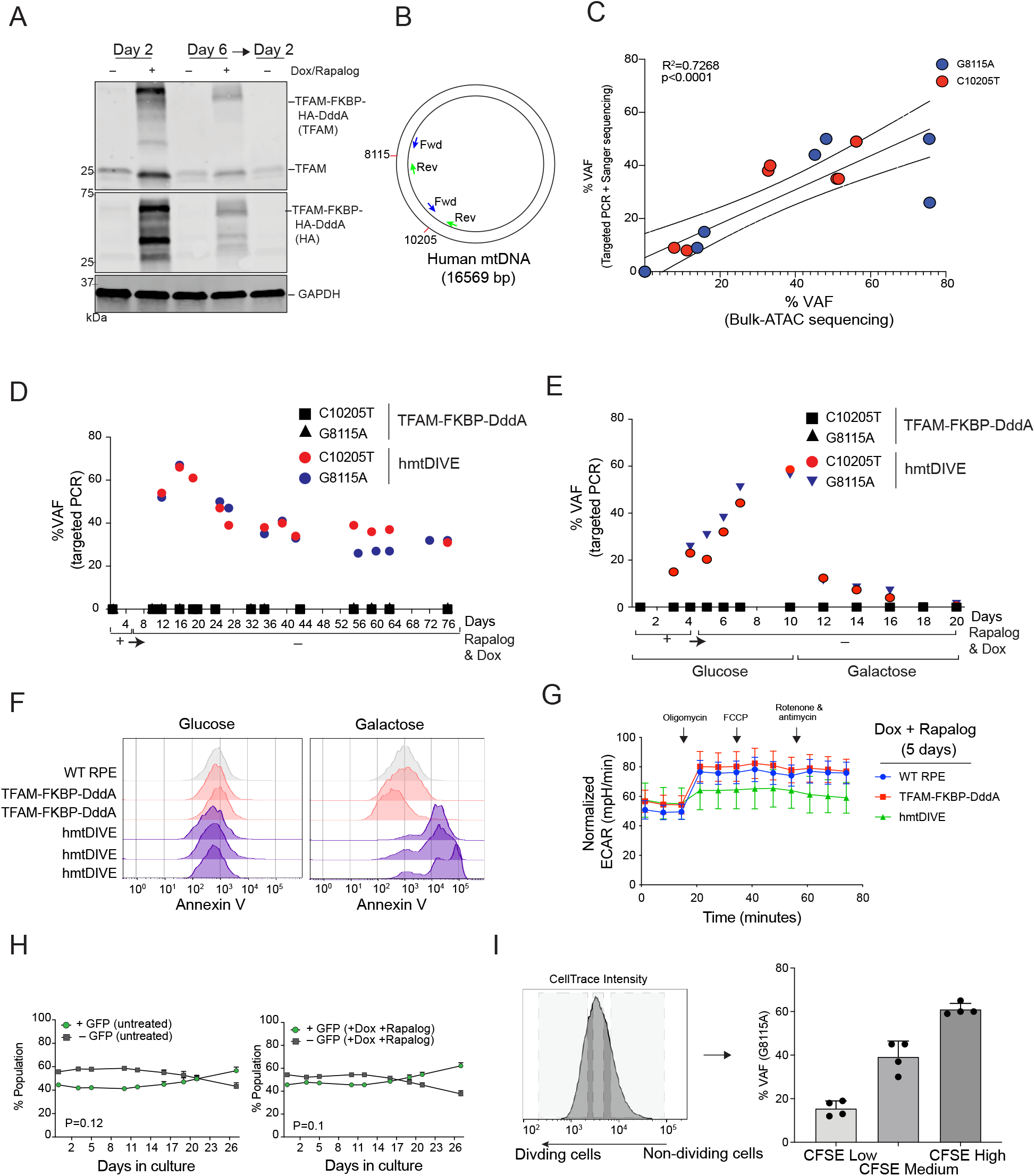
Metabolic context dictates the mode of selection against mtDNA mutations. **(A)** Western blot analysis of the doxycycline-inducible HA-tagged FKBP-TFAM-DddA fragment in cells after 2 days and 6 days of doxycycline and rapalog treatment. TFAM and HA immunoblots are shown; GAPDH serves as a loading control. **(B)** Circular map of the human mitochondrial genome showing primer positions used for targeted amplification of two recurrent mtDNA variants: G8115A (MT-COX2) and C10205T (MT-ND4). Forward (Fwd) and reverse (Rev) primer positions are indicated. **(C)** Correlation between VAF measurements obtained by targeted PCR followed by Sanger sequencing and bulk ATAC-seq. Data points represent measurements of C10205T (red) and G8115A (blue) across samples from one biological replicate. Lines indicate the linear regression fit and 95% confidence interval. **(D)** Longitudinal tracking of C10205T and G8115A VAF over 76 days. Cells were treated with doxycycline and rapalog during an induction period (days 0–5), then cultured without drugs (days 6–76). TFAM-FKBP-DddA control cells are shown in black; hmtDIVE-expressing cells in red and blue. **(E)** Time-course measurement of C10205T and G8115A VAF under glucose and galactose conditions in cells expressing TFAM-FKBP-DddA (black) and hmtDIVE (red/blue). Doxycycline and rapalog were administered during the induction period (days 0-5) and withdrawn during days 6-20. (D-E) VAF was measured by Sanger sequencing as described in the Methods. **(F)** Annexin V staining by flow cytometry to assess apoptotic cell fractions in WT RPE, TFAM-FKBP-DddA, and hmtDIVE-expressing cells cultured in glucose or galactose for 2 days. Overlay histograms show Annexin V fluorescence intensity across biological replicates. **(G)** Extracellular acidification rate (ECAR) measurements in WT RPE (blue), TFAM-FKBP-DddA (red), and hmtDIVE-expressing (green) cells after 5 days of doxycycline and rapalog induction. ECAR was measured following sequential addition of oligomycin, FCCP, and rotenone + antimycin. Values were normalized to cell number; data represent mean ± SD from *n* = 3 biological replicates. **(H)** Population dynamics of hmtDIVE-expressing cells tracked over 26 days, with both GFP^+^ and GFP^-^ cells receiving the same treatment. Left: untreated GFP^+^ (green) and GFP^-^ (gray) hmtDIVE-expressing cells. Right: GFP^+^ (green) and GFP^-^ (gray) hmtDIVE cells treated with doxycycline and rapalog. The relative abundance of GFP^+^ cells was quantified by flow cytometry over time. Data represent mean ± SD from *n =* 3 biological replicates. P-values were derived from a two-sided t-test of the normalized area under the change-from-baseline curve against zero. **(I)** CellTrace proliferation analysis of hmtDIVE-expressing cells. Left: representative histogram showing CFSE fluorescence used to define divided and undivided cell populations. Right: G8115A VAF in CFSE-low (divided), medium, and high (undivided) populations. Data represent mean ± SD from *n =* 4 biological replicates.

**Figure S3:**
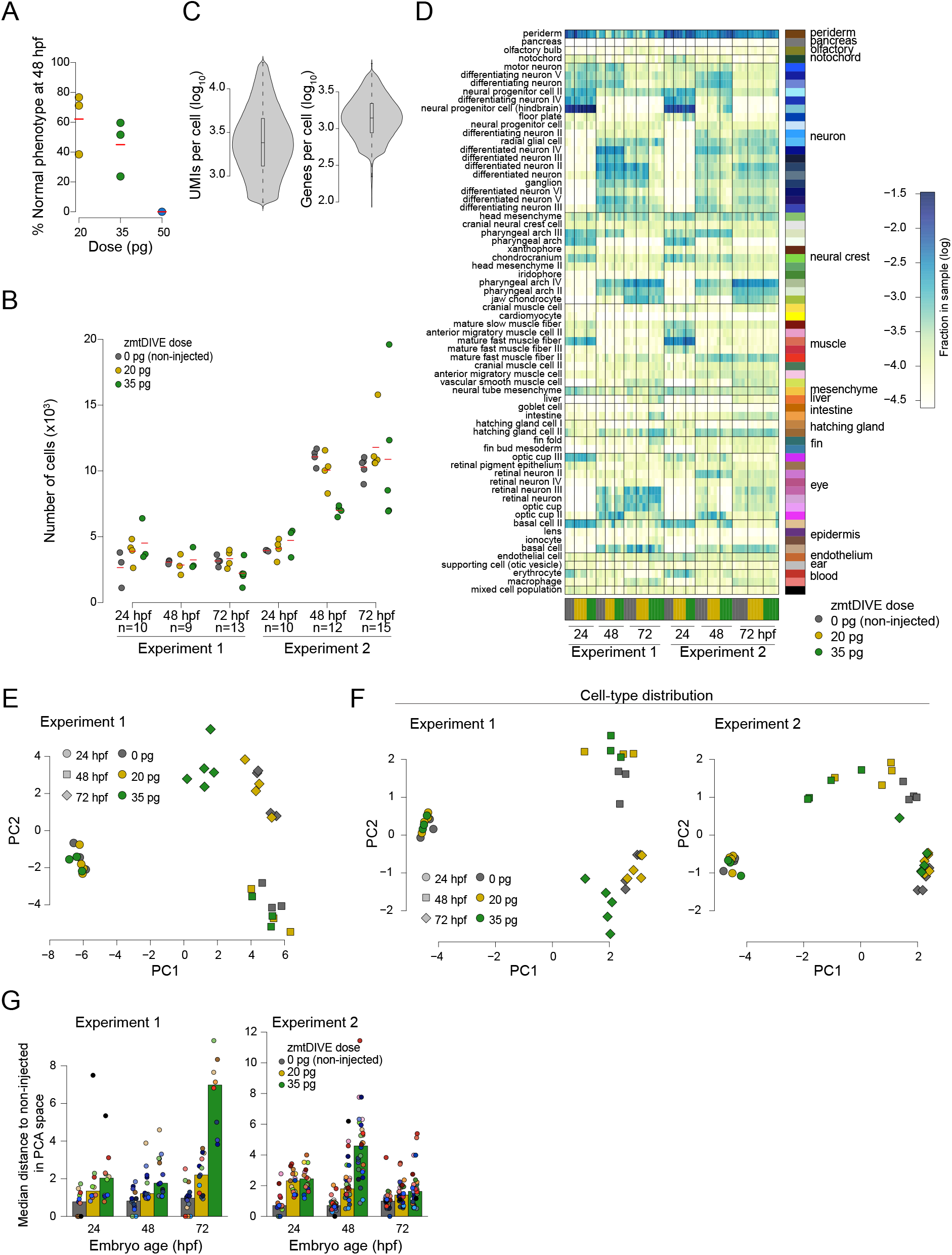
Global transcriptional effect of zmtDIVE on early development. **(A)** Fraction of morphologically-normal zmtDIVE-injected embryos across varying doses at 48 hpf. Each dot represents an independent batch of injected embryos. N=3 (20 pg, 35 pg), 1 (50 pg) injection batches. Each batch consists of 38-72 injected embryos. **(B)** Number of recovered cells per embryo across two independent experiments, three time points, and three zmtDIVE doses. **(C)** Distribution of UMI counts and number of detected genes per cell across all profiled single cells. **(D)** Distribution of cell types across all profiled embryos. The matrix is column-normalized and log-transformed. **(E)** Principal Component Analysis (PCA) of pooled gene expression profiles from 32 injected embryos in Experiment 1. Each symbol represents one embryo. Symbol shape indicates embryo age and color indicates zmtDIVE dose. **(F)** Similar to (E), but computed using normalized cell type distributions. **(G)** Distance in PCA space between each cell type and age-matched non-injected controls. Bars represent the mean distance across all cell types. Each dot represents one cell type; cells were pooled from embryos of the same developmental stage and injection dose.

**Figure S4:**
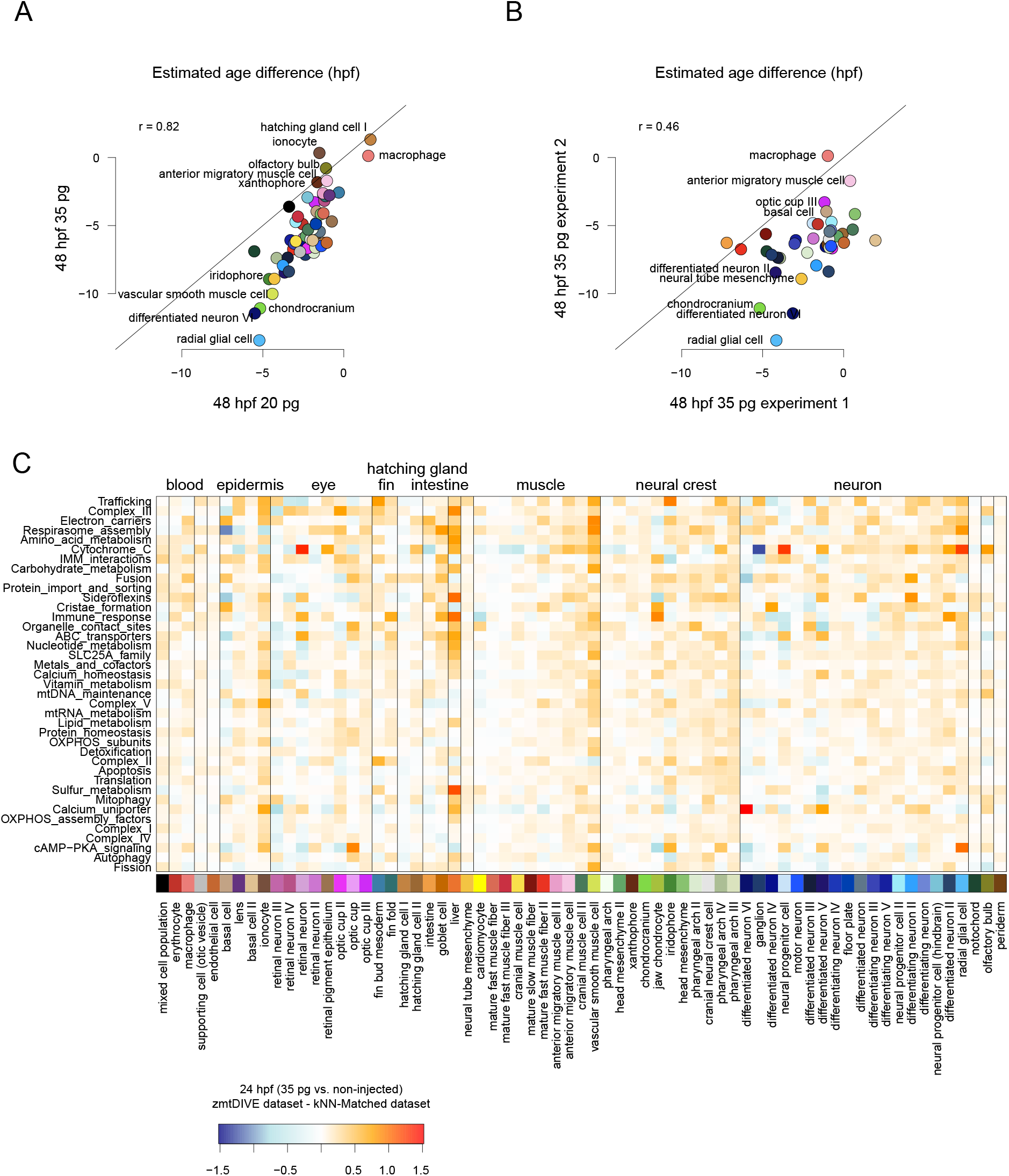
Transcriptional and developmental effects of zmtDIVE injection. **(A-B)** Difference in estimated developmental age across cell types. Each dot represents one cell type. **(A)** Comparison of 48 hpf embryos injected with different RNA doses. **(B)** Comparison of 48 hpf embryos injected with 35 pg zmtDIVE in two independent experiments. **(C)** Enrichment (log_2_ fold change) of nuclear-encoded mitochondrial gene modules (rows)compared to non-injected controls across cell types of 35 pg-injected embryos at 24 hpf (columns). Values are normalized to the enrichment calculated between stage-matched reference cells corresponding to the k-nearest neighbors (K = 10) of 35 pg–injected and non-injected embryos in the developmental atlas, thereby negating the effect of developmental delay.

**Figure S5:**
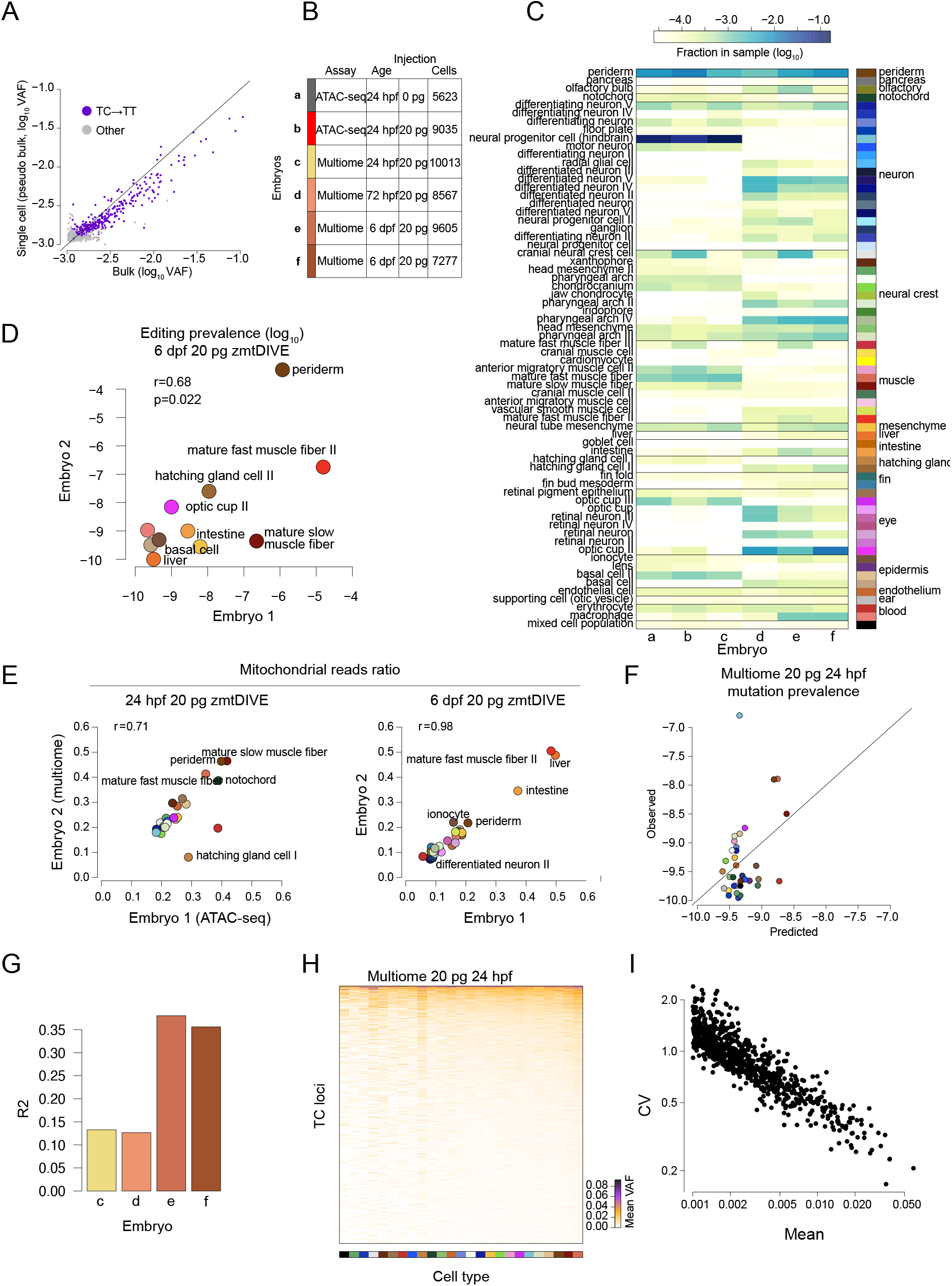
Cell-type differences in zmtDIVE-induced mutation prevalence. **(A)** Comparison of C→T VAF (log_10_) between two 20 pg zmtDIVE-injected embryos at 24 hpf. The x-axis shows VAF from bulk sequencing (as in **Fig. S1E**), and the y-axis shows pooled reads from a single-cell mt-enriched ATAC-seq sample, demonstrating concordance across platforms. Purple dots indicate TC motifs. Only loci with sufficient coverage in both samples are shown. **(B)** Summary of six mt-enriched single-cell ATAC-seq and multiome samples from zmtDIVE embryos. **(C)** Cell-type composition across the six profiled embryo samples. Columns represent embryos as in (B). **(D)** Pearson correlation of average mtDNA mutation prevalence across cell types between two 20 pg zmtDIVE-injected embryos 6 dpf. Each point represents a cell type. Values represent the geometric mean fraction of cells carrying a mutation across all TC motifs. **(E)** Pearson correlation of the fraction of mitochondria-aligned reads across cell types between biological replicates. Left: comparison between an ATAC-seq and multiome dataset at 24 hpf. Right: comparison between two 6 dpf replicates. Each point represents a cell type. **(F-G)** Performance of a multivariable model predicting cel-type-specific zmtDIVE mutation prevalence from measured cell type features. **(F)** Observed versus predicted editing prevalence across cell types for a representative embryo (sample c in Fig. S5B). The model incorporates fraction of mitochondrial reads (as in (E)), total UMI count, and fraction of mitochondrial transcripts. **(G)** Model performance, measured as *R*^2^, across embryos. Columns represent embryos as in (B). **(H-I)** Cell type-specific zmtDIVE activity across loci. **(H)** Mean VAF across loci (rows) and cell types (columns) after column-wise L2 normalization. **(I)** Coefficient of variation (CV) as a function of mean editing level, indicating uniform scaling across loci. Values are log-transformed.

**Figure S6:**
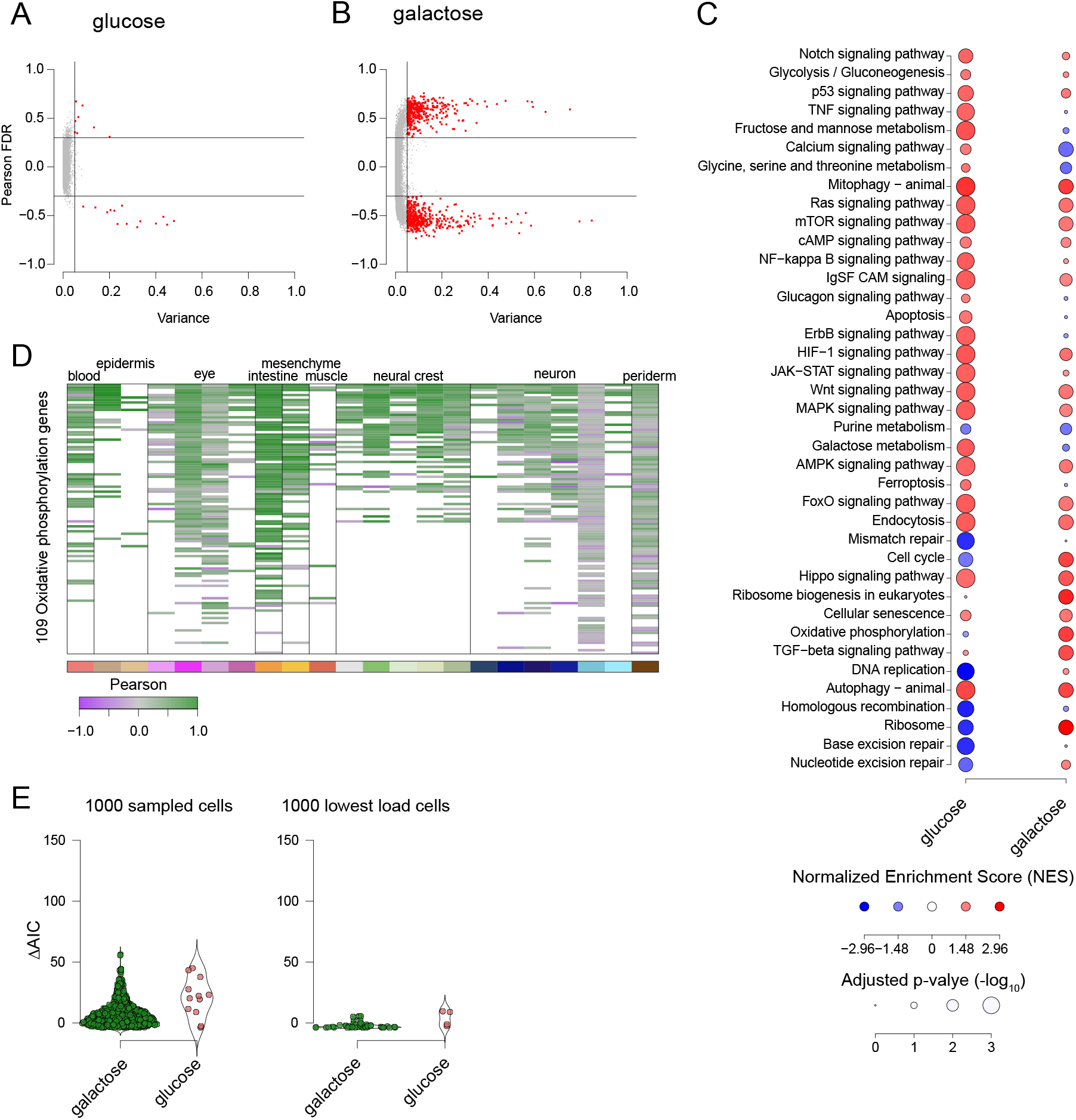
Transcriptional response to elevated mutation load. **(A-B)** Pearson correlation between gene expression and mutation load plotted against expression variance in cells cultured for 1 day in glucose (A) or galactose (B). Each dot represents a gene. Genes with r>0.3 and variance>0.05 are highlighted in red. Correlation and variance were calculated across bins of cells (k=30) stratified by mutation load. **(C)** Gene Set Enrichment Analysis (GSEA) of gene programs associated with mutation load in hmtDIVE-expressing cells cultured for 1 day in glucose or galactose. Circle color indicates normalized enrichment score (NES); circle size reflects statistical significance (FDR-adjusted p-value). **(D)** Heatmap showing Pearson correlations between mutation load and expression of 109 genes belonging to the oxidative phosphorylation KEGG annotation across 22 cell types, grouped by developmental lineage. Missing values indicate genes with insufficiently variable expression. **(E)** Distribution of ΔAIC values across hmtDIVE-induced cells cultured for 1 day in glucose or galactose as in Fig. 5D. Shown are ΔAIC values calculated after down-sampling cells from each condition to 1,000 cells (left), or after selecting the 1,000 cells with the lowest mutation load (right). Each dot represents a load-responsive gene. Positive ΔAIC values indicate greater support for threshold-like responses, whereas negative values indicate greater support for linear responses.

**Figure S7:**
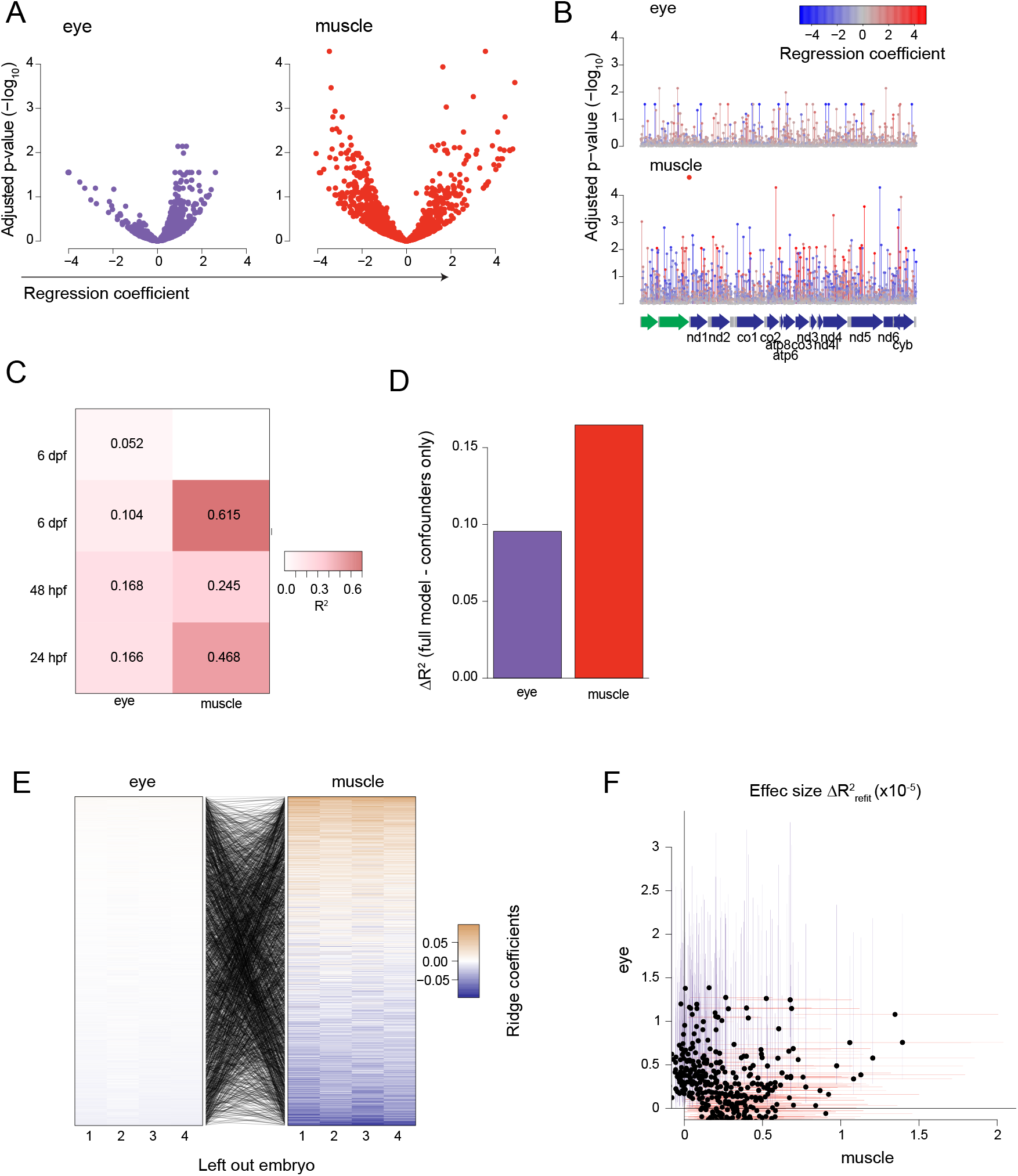
Individual mtDNA mutations differentially contribute to cellular response across zebrafish lineages. **(A)** Linear regression analysis estimating the contribution of individual loci to a gene score representing the oxidative phosphorylation program across two developmental lineages (eye and muscle). Models control for embryo, fraction of mtDNA reads, and global mutation load excluding the tested locus. Each dot is a TC mtDNA locus. Values indicate FDR-adjusted p-values (y-axis) plotted against regression coefficients (x-axis). **(B)** Lollipop plots showing the statistical significance (-log_10_ FDR-adjusted p-value) of individual loci in the two models in (A), across the mitochondrial genome. Locus color indicates the corresponding model coefficient. Bottom: mitochondrial genome maps positioning variants relative to mtDNA-encoded genes. Protein-coding genes are shown in blue, rRNA in green and tRNA genes in gray. **(C)** Performance of lineage-specific cross-validated ridge regression models across four embryos. Values indicate R^2^ on held-out test sets. Missing values indicate an insufficient number of cells in the test set. **(D)** Additional explained variance attributable to individual mtDNA loci in two lineage-specific models. Values represent ΔR^2^, calculated as the difference in R^2^ between the model incorporating full mutation profiles and a minimal model containing global mutation load and technical covariates only. **(E)** Robustness of mtDNA regression coefficients across leave-one-embryo-out refits. Lines between heatmaps connect coefficients for the same loci across cell lineages. **(F)** Comparison of locus-specific effect sizes between muscle (x axis) and eye (y axis) ridge regression models. Effect size was calculated as the decrease in model performance ΔR^2^(refit) after removing a locus and refitting the model. Each dot represents a locus; colored lines indicate 95% confidence intervals across 500 bootstrap iterations.

**Figure S8:**
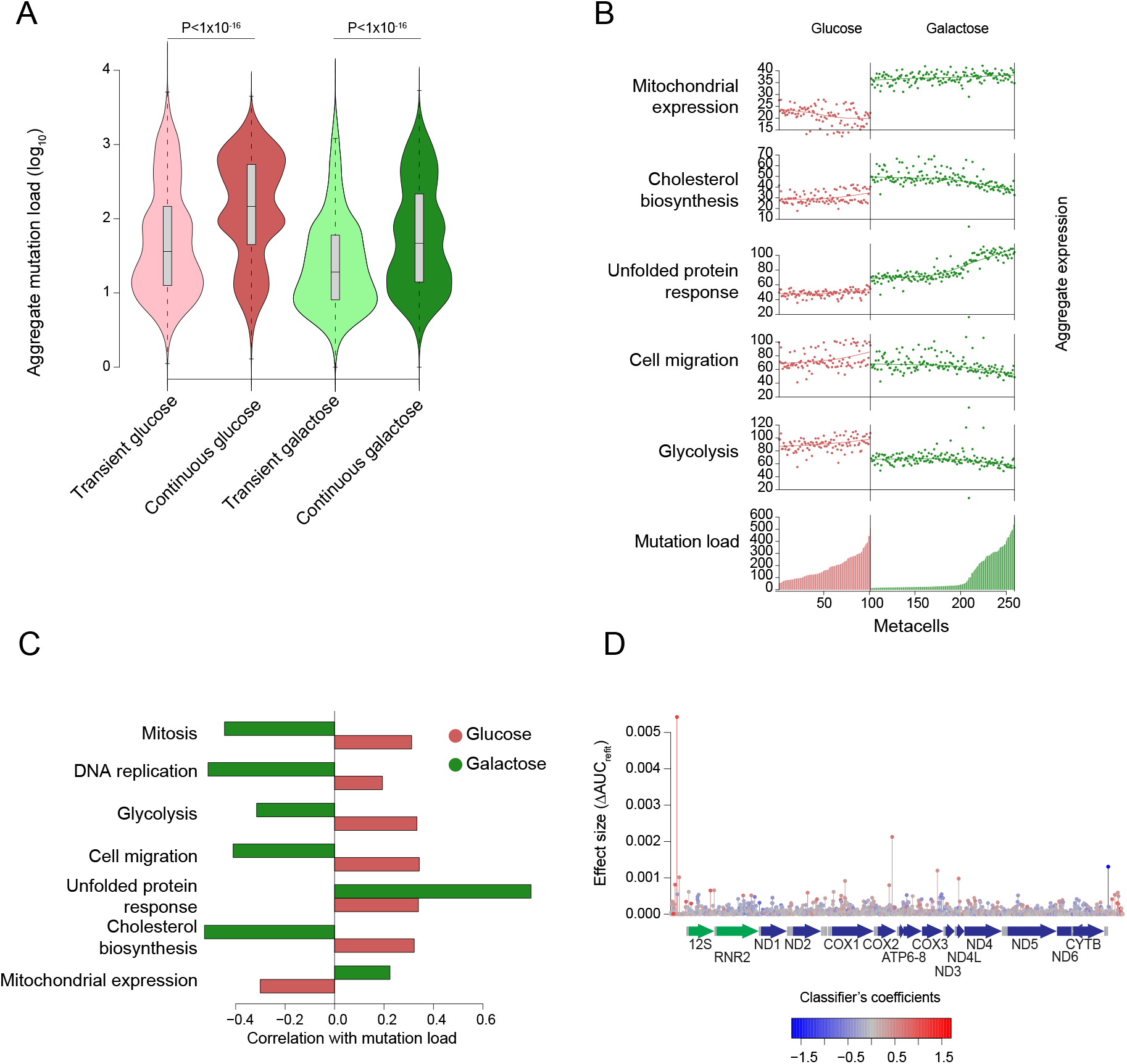
Contrasting mitochondrial transcription under constitutive hmtDIVE expression. **(A)** Distribution of aggregated mutation load across cells cultured for 10 days in glucose or galactose under transient or continuous hmtDIVE induction. Mutation load was calculated as the sum of the log Bayes factors across all TC loci in the mitochondrial genome. Two-tailed unpaired Mann-Whitney test. **(B)** Expression of gene expression programs across metacells from cells grown for 10 days under continuous hmtDIVE induction in glucose or galactose. Metacells are ordered by median aggregated mutation load. **(C)** Pearson correlation between gene-program expression and mutation load across metacells as in (B), relating to glucose or galactose growth conditions. **(D)** Lollipop plots showing the effect sizes of individual loci as in Fig. 6E, plotted across the mitochondrial genome. Locus color indicates the classifier’s coefficient. Bottom: mitochondrial genome maps positioning variants relative to mtDNA-encoded genes. Protein-coding genes are shown in blue, rRNA in green and tRNA genes in gray.

**Figure S9:**
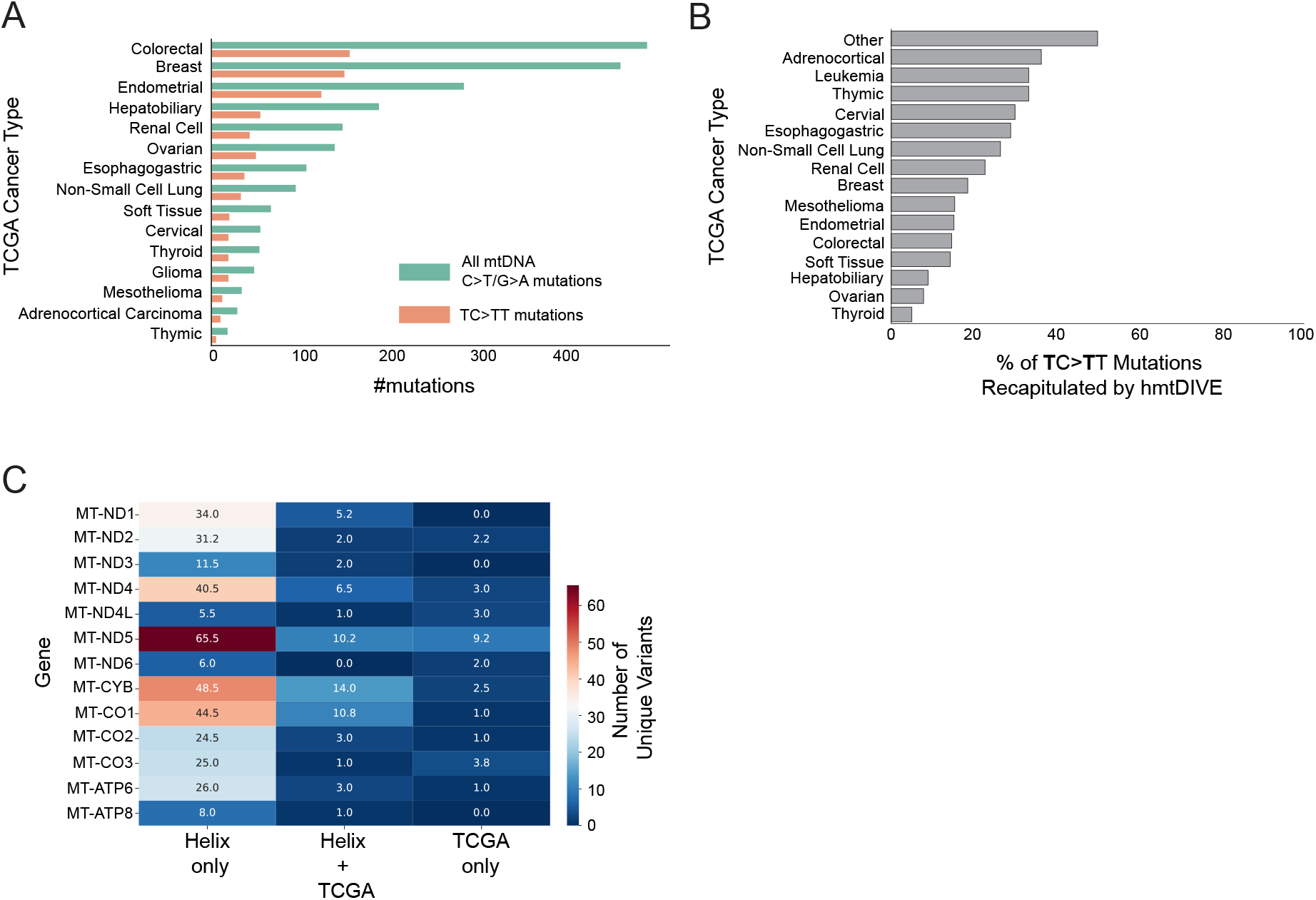
Representation of hmtDIVE-induced mutations in cancer and population polymorphism. **(A)** Number of mtDNA C→T mutations and motif-specific TC→TT mutations identified in TCGA tumor samples, stratified by cancer type. **(B)** Percentage of motif-specific mtDNA mutations (TC→TT) identified in each TCGA cancer type that are recapitulated by hmtDIVE. Percentages are calculated relative to the total number of motif-specific variants per cancer type. **(C)** Heatmap showing the number of unique TC→TT variants per gene, categorized by their presence in Helix-only, shared (Helix + TCGA), or tumor-only (TCGA exclusive) datasets. Each cell represents the mean number of unique variants per category across *n =* 4 biological replicates. Color reflects the number of unique variants.

## SUPPLEMENTARY TABLES

Table S1: Average gene expression profiles of key marker genes across 13,010 metacells. Metacells are annotated into cell types, and genes are grouped into annotation groups based on Saunders et al.

Table S2: Summary of hmtDIVE samples sequenced by DOGMA-seq

Table S3: Grouping of zebrafish cell types into developmental lineages

Table S4: Per-locus summary of hmtDIVE mitochondrial-activity classifier coefficients, effect sizes, and mutation metadata.

Table S5: Primers used to amplify regions in zebrafish mtDNA

